# Opening the species box: What parsimonious microscopic models of speciation have to say about macroevolution

**DOI:** 10.1101/2023.11.09.564915

**Authors:** Élisa Couvert, François Bienvenu, Jean-Jil Duchamps, Adélie Erard, Verónica Miró Pina, Emmanuel Schertzer, Amaury Lambert

**Affiliations:** Institut de Biologie de l’ENS (IBENS), École Normale Supérieure, PSL Université, CNRS UMR8197, INSERM U1024, Paris, France; Centre Interdisciplinaire de Recherche en Biologie (CIRB), Collège de France, PSL Université, CNRS UMR7241, INSERM U1050, Paris, France; Université Paris Cité, CNRS UMR8145, MAP5, F-75006, Paris, France; Université de Franche-Comté, CNRS, LmB (UMR 6623), F-25000 Besançon, France; Centre for Genomic Regulation (CRG), The Barcelona Institute of Science and Technology, Barcelona, Spain; Universitat Pompeu Fabra (UPF), Barcelona, Spain; Faculty of Mathematics, University of Vienna, Oskar-Morgenstern-Platz 1, 1090 Wien, Austria

## Abstract

In the last two decades, lineage-based models of diversification, where species are viewed as particles that can divide (speciate) or die (become extinct) at rates depending on some evolving trait, have been very popular tools to study macroevolutionary processes. Here, we argue that this approach cannot be used to break down the inner workings of species diversification and that “opening the species box” is necessary to understand the causes of macroevolution, but that too detailed speciation models also fail to make robust macroevolutionary predictions.

We set up a general framework for parsimonious models of speciation that rely on a minimal number of mechanistic principles: (i) reproductive isolation is caused by excessive dissimilarity between genotypes; (ii) dissimilarity results from a balance between differentiation processes and homogenizing processes; and (iii) dissimilarity can feed back on these processes by decelerating homogenization.

We classify such models according to the main homogenizing process : (1) clonal evolution models (ecological drift), (2) models of genetic isolation (gene flow) and (3) models of isolation by distance (spatial drift). We review these models and their specific predictions on macroscopic variables such as species abundances, speciation rates, interfertility relationships or phylogenetic tree structure.

We propose new avenues of research by displaying conceptual questions remaining to be solved and new models to address them: the failure of speciation at secondary contact, the feedback of dissimilarity on homogenization, the emergence in space of breeding barriers.

## 1 Introduction

### 1.1 Phylogenetic approaches to diversification: let us open the species box

Starting with the seminal works of the “Woods Hole group” paleontologists (Raup et al., 1973) and drawing on parallel mathematical progress (Kendall, 1948; Nee et al., 1994; Aldous, 2001; Aldous and Popovic, 2005), a powerful quantitative method has been developed in macroevolution, using birth-death processes as models for species diversification. In these so-called *lineage-based models of diversification*, species are particles that can undergo two kinds of events: speciation, modeled by instantaneous division; and extinction, modeled by instantaneous death. The phylogenetic patterns (as quantified by tree balance indices and other tree shape statistics, see Box 4) predicted by lineage-based models can then be studied mathematically and used either to test whether a birth-death process, seen as a null model, can explain the observed phylogeny; or, alternatively, to infer how speciation and extinction rates may depend on some evolving trait carried by the species. This so-called *phylogenetic approach to diversification* has been very popular in macroevolution and surveyed multiple times in the last decade (Ricklefs, 2007; Pyron and Burbrink, 2013; Stadler, 2013; Morlon, 2014; Morlon et al., 2024).

However, when it comes to processes as complex – and occurring on time scales as long – as species diversification, this approach suffers from several limitations:

- The build-up of genetic differentiation between populations that leads to the formation of new species takes time, and so do the demographic declines and population extirpations that lead to species extinctions. This is not captured by coarse-grained lineage-based models where speciation and extinction are instantaneous;
- A single phylogeny contains little signal, which gives rise to statistical problems like false associations between rate and trait (Rabosky and Goldberg, 2015) or non-identifiability of parameters (Louca and Pennell, 2020) – but see also Morlon et al. (2022) for a discussion;
- Phylogenies are nowadays built by comparing sequences of nucleic acids, hence the name “molecular phylogenies”. However, because of recombination and, as increasingly recognized, gene flow between species (Marques et al., 2019; Pennisi, 2016), different genes can have very different genealogies, so that a phylogeny is merely a brief summary of evolutionary history (Degnan and Rosenberg, 2006; Maddison, 1997);
- Speciation and extinction rates are useful notions to understand the diversification process but ultimately, are only compounded quantities that summarize very crudely some finescale phenomena such as habitat selection, species sorting, divergent adaptation, reproductive isolation, assortative mating, introgression, reinforcement, speciation collapse, etc. If we want to characterize the determinants of species diversification, we have to understand and infer these processes (Li et al., 2018; Rolland et al., 2023; Morlon et al., 2024; Harvey et al., 2017, 2019). Actually, the way diversification rates and other macroevolutionary or macroecological observables depend on these fine-scale processes remains one of the most intriguing questions in macroevolution.

To address these questions, an alternative way to study diversification processes consists in moving away from the assumption that species are particles (top-down approach) to directly model how species’ elementary constituents (populations, individuals, genomes) all work together to lead to speciation (bottom-up approach). Models pertaining to the bottom-up approach are rooted in the fine-scale description of ecological and genetic phenomena and called **microscopic models**.

In the next section, we argue that microscopic models need to remain parsimonious to make robust macroscopic predictions and we explain how to conceive and study such models.

### 1.2 A plea for keeping microscopic models simple

In evolutionary biology, a large and long-standing body of theory seeks to model the process of speciation in order to go beyond verbal theories and quantify the effects of the various mechanisms at work (Coyne et al., 2004; Turelli et al., 2001). The first such models used the framework of population genetics to address the evolution of postzygotic isolation and date back to Dobzhansky (1937); Wright (1941); Muller (1942). Subsequent works, in particular those seeking to study prezygotic isolation, also model ecological processes such as mating and dispersal. A growing part of speciation theory has then left mathematical analysis to the benefit of numerical simulations, allowing models to become more and more complex. For example, individual-based models of ecological speciation may rely on a detailed description of the ecology where each individual is identified with a quantitative trait (e.g., foraging strategy), a mating trait (trait driving mating preferences) and sometimes even a spatial position (e.g., Dieckmann and Doebeli, 1999; Doebeli and Dieckmann, 2003; Thibert-plante and Hendry, 2009; Aguilée et al., 2011, 2013; Gascuel et al., 2015; Aguilée et al., 2018).

Such parameter-rich models sacrifice simplicity for precision and realism. Their predictions are always suspect of depending on specific modeling choices and of being valid only in some regions of parameter space (Turelli et al., 2001). In terms of parameter inference, such models can easily suffer from over-fitting and/or non-identifiability which makes difficult to relate reliably microscopic parameters and macroevolutionary observables.

Therefore, we argue in favor of a balance between top-down and bottom-up approaches: while definitely needing to open the species box, at the same time must we focus, to identify the drivers of macroevolution, on microscopic models that remain parsimonious.

We use the term **archetypal** to refer to models that are both **microscopic** and **parsimonious**. Building and using archetypal models is a 4-step process drawing on complementary tools and areas of expertise. Hereafter, we give a brief description of these four steps and of their benefits in the broader field of population biology.

First, microscopic models are specified by relying on **mechanistic principles** and **measurable parameters** (e.g., dispersal rate, demographic parameters, mutation rate). One way of keeping the number of parameters low is to overlook selective processes or to model them in a nonparametric way using e.g., effective or composite parameters, holey landscapes, rank-based selection.

As a second step, **mathematical analyses** can provide an assessment of the range of application of the model, also called its “class of universality”. In its weak sense, universality here refers to formalisms that can fit a wide variety of phenomena, because they heuristically picture what are suspected to be the main mechanisms at work or because several mechanisms can be represented by the same formalism.

In its strong sense, universality refers to mathematical micro–macro approaches where a large class of models can be approximated by a limit model with few parameters, sometimes at the cost of taking parameters of the initial models to some extremal region of the parameter space (e.g., unlimited dispersal, also called mean field limit; large community size; small mutation rate; etc.). This approach is standard in population genetics, where stochastic models with explicit genealogy and constant population size *N* (Cannings models) with neutral mutation rate *u_N_* converge as *Nu_N_ θ* towards a universal genealogy (Kingman coalescent) with Poissonian mutations with intensity *θ*. In between these two extremes lie models using composite parameters that can summarize a limited range of mechanisms.

The third step then consists in deriving accurate predictions of the model for some **macroscopic observables** corresponding to biological quantities of interest, either at the species level (intraspecific genetic diversity, species abundance, species range) or at the community level (species richness, speciation/extinction rate, phylogeny). These macroscopic observables can also be more complex patterns, such as relations between two kinds of observables (species-area relationship, species-speciation rate relationship, ecosystem functioning-diversity relationship, gene tree-species tree coupling).

The last step consists in confronting the model to reality. This involves tuning parameters to see how well the model’s predictions can fit the data (typically, using maximum likelihood estimates).

When successful – which requires that the mathematical analysis work and that the predictions fit the data –, this methodology produces models that

1. Are biologically realistic (step 1);
2. Are universal, in the sense that they reflect the general behavior of a large class of more detailed models (step 2);
3. Yield predictions that can readily be linked back to the basic assumptions (step 3);
4. Have few parameters and therefore are easily falsifiable, which gives more confidence in their explanatory power (step 4).

We explain how this generic strategy can be instantiated for speciation research when introducing archetypal models of speciation in forthcoming Section 1.4 and in some dedicated boxes.

### 1.3 The species definition problem

The bottom-up approach to speciation gives insight into what makes a species a species. Indeed, despite being central to evolutionary biology, the notion of species is not well-defined, in the sense that there is no single agreed formal definition. One of the most widely accepted ideas is that species should correspond to “groups of reproductively compatible populations that are reproductively incompatible with other such groups”. This is the so-called *biological species concept*, or BSC; see Coyne et al. (2000) for a detailed discussion.

However, despite its apparent simplicity – and considerable influence in biology – the BSC admits more than one mathematical formalization (see Box 1). More than a problem with the BSC, this reflects the fact that species are complex entities with fuzzy boundaries. Much like when trying to define a heap of sand, it may not be possible to find a one-size-fits-all definition. But this does not prevent dunes from existing, nor does it make it impossible to study their formation and dynamics. Only by opening the species box and by choosing the right level of description to model species’ elementary constituents – metapopulations, populations, individuals, genes – can we hope to circumvent the “species problem” and to model species formation without being tied to a specific (and inevitably imperfect) definition.

#### Box 1

##### The biological species concept and the interfertility graph.

The *biological species concept* (BSC) considers species to be “groups of reproductively compatible populations that are reproductively incompatible with other such groups”. If this were a strict definition, then phenomena such as ring species (where populations of the same species are reproductively incompatible, see Irwin et al. 2001) and hybrid speciation (where populations of different species hybridize to produce a new species, see Mallet 2007) would be impossible. Therefore, the BSC tells us what the essence of an idealized species should be, but it is not an accurate description of species in the real world.

The BSC framework postulates that the key criterion on which species should be defined is the notion of *interfertility*, i.e. the ability to interbreed. Interfertility is, ultimately, a relation between individuals; but as a first step in opening the species box it can be seen as a relation between populations. This relation can be represented by a graph whose vertices correspond to populations, and where two vertices are linked by an edge if and only if the two corresponding populations are reproductively compatible: we call this graph the *interfertility graph*. In an ideal setting, the interfertility graph would be a disjoint union of cliques (i.e. groups of vertices such that there is an edge between every pair of vertices), but – as we have seen – it will deviate from this ideal in practice. Bienvenu et al. (2019) introduced a mathematical model to study the structure and dynamics of interfertility graphs. The following graphs are simulations from that model, for 1000 populations and various values of its drift parameter.

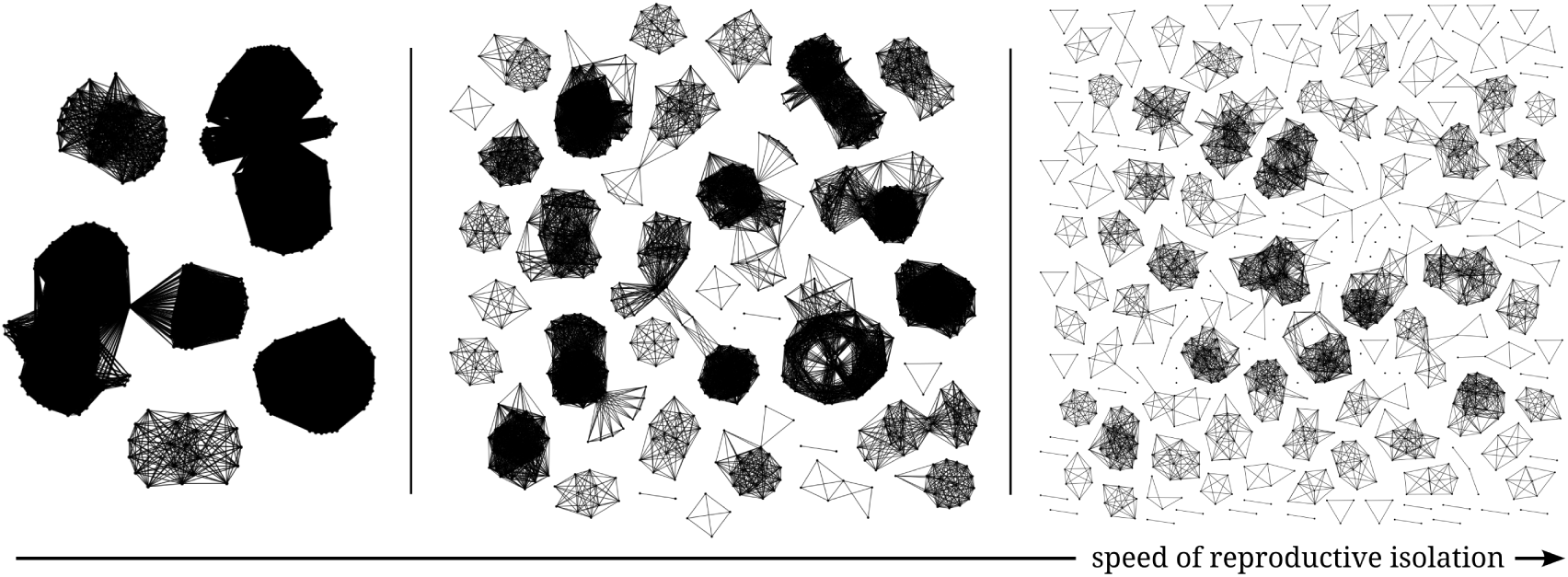

Given their interfertility graph, there is a continuum of ways to partition a set of populations into species, as illustrated in Figure 1. At one end of the spectrum, if we want to ensure that reproductively compatible populations belong to the same species, then species have to be defined as maximal cliques; however, some drawbacks of this approach are that there may not be a unique partition into maximal cliques, and that it may allow too much hybridization. At the other end of the spectrum, species can be defined as connected components of the interfertility graph. This definition has the advantage of being unambiguous and very natural, in that two populations belong to the same species if and only if there is some possibility of (direct or indirect) gene flow between them; however, it can lead to species being composed mostly of reproductively incompatible populations, and it does not allow any hybridization. Thus, there is no “one-size-fits-all” definition, and which definition proves most relevant may depend on the specific setting.

Last, note that in reality interfertility may vary along the genome, for example when an inversion prevents parts of the genome to get exchanged, or when disruptive selection blocks gene flow across blocks of the genome linked to causal loci. See Wu (2001) for a more detailed discussion on this topic.

**Figure 1:**
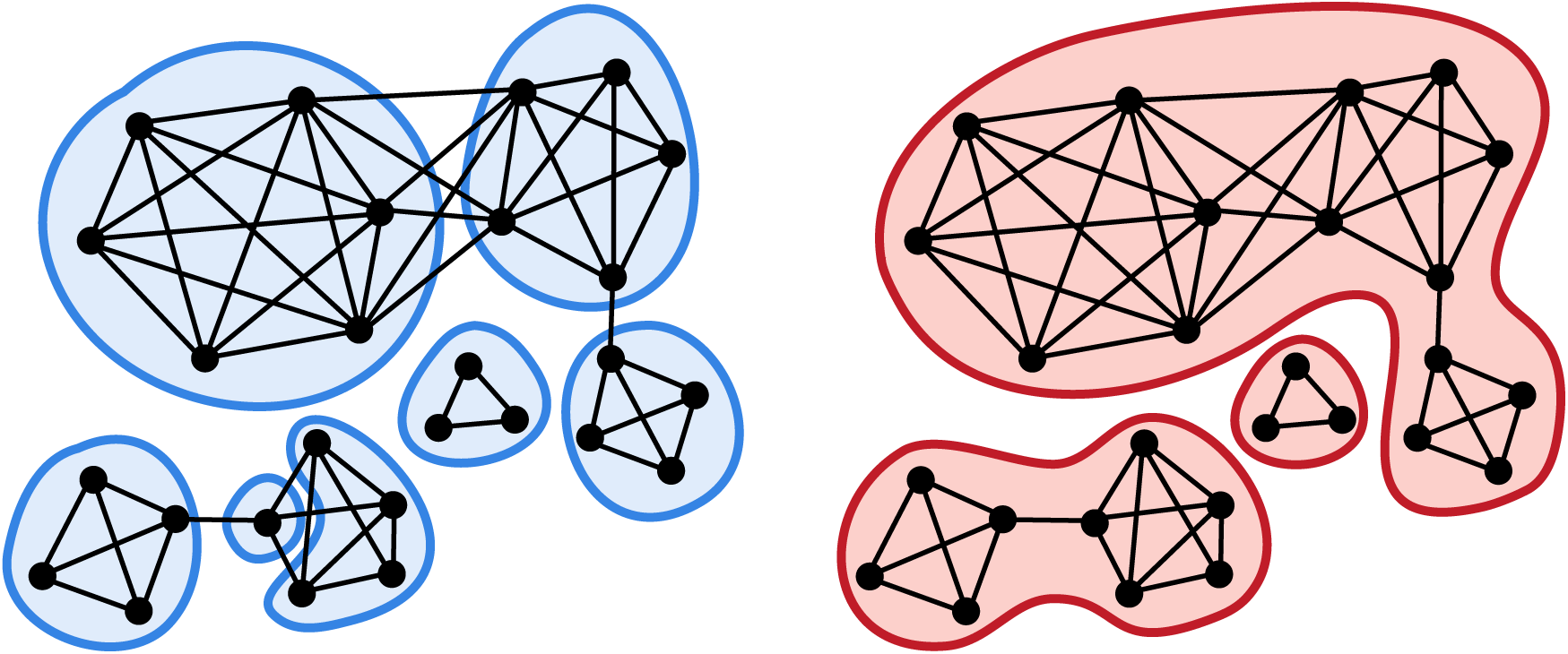
Several ways to partition the vertices of a graph. On the left, in blue, a partition into maximal cliques. On the right, in red, the partition into connected components.

On the other hand, for any microscopic model representing the speciation process, one has to decide, after letting the process unfold, which groups of populations or individuals have to be considered species, in order to uncover the model’s macroecological predictions. This procedure, which consists in assigning each individual/population of a sample to a species, is known as the *species clustering problem*. This problem is not only relevant to our theoretical understanding of speciation: it also arises as a very concrete question in microbial genomics and metagenomics, particularly in the context of barcoding and metabarcoding.

In view of the BSC, a natural way to group individuals/populations into species is to consider all relations of interfertility and to partition the associated interfertility graph into groups such that each pair of elements in the same group are linked by a chain of interbreeding pairs, see Box 1. Figure 1 displays two such partitions: into maximal cliques or into connected components.

However in most cases, the interfertility graph is not available and one has to resort to more easily accessible kinds of data – typically genetic distances. Given a matrix of genetic distances between individuals, the following algorithm, which we refer to as the *threshold clustering algorithm*, provides a solution to the species clustering problem:

- Fix a threshold distance;
- Consider the graph where any two individuals are linked if and only if the genetic distance between them does not exceed this threshold;
- Species are then defined to be the connected components of this graph.

For example, metabarcoding studies traditionally use a threshold clustering algorithm to delimit species (called “operational taxonomic units” or OTUs in that context), using the percentage of pairwise differences on 16S ribosomal RNA for the genetic distance, and a threshold equal to 0.03.

Many microscopic models of speciation implicitly use a threshold clustering algorithm to define species. In most cases, lineages are assumed to accumulate differences under the *infinite-allele model* (Kimura and Crow, 1964). In this setting, each mutational event gives rise to a new allele that has never existed in the past and increases the genetic distance to each ancestor by 1. More formally, the infinite-allele assumption endows the alleles with a tree structure called the *allelic tree*, see Figure 2: allele *A* is the mother of allele *B* if the mutation giving rise to allele *B* occurred on a lineage carrying allele *A*. The genetic distance between two individuals is merely the graph distance between their alleles in the allelic tree. In the case when the threshold is taken to be 1, the species partition thus obtained is the finest partition such that two individuals carrying the same allele are found in the same species.

**Figure 2:**
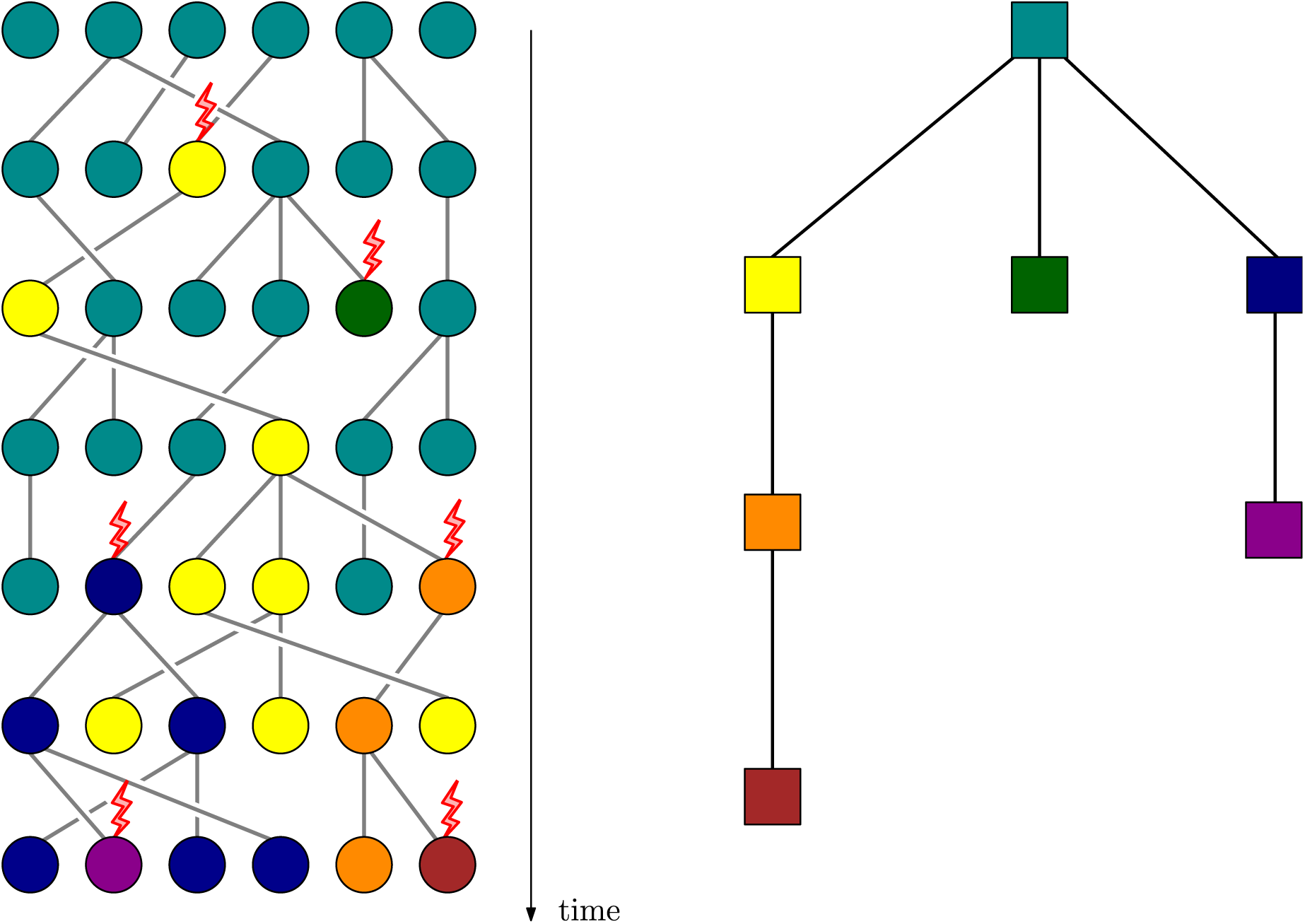
An individual-based genealogical model with mutations under the infinite-allele model. Left: each circle represents an individual, and each color an allelic type. A red lightning indicates a mutation, yielding a new type. Right: the allelic tree corresponding to the process on the left.

Appealing though they may be for their elegance and practicality, a major pitfall of threshold clustering algorithms is that, even under simple models of evolution of genetic distances, they do not always yield species partitions that are compatible with the genealogy – that is, the corresponding “species” are not always monophyletic (because, for example, of incomplete lineage sorting or hybrid speciation; see also Figure 1 of Manceau and Lambert, 2019). This raises the fundamental question of the existence of a natural species partition such that (1) each species forms a subtree of the genealogy and (2) any two individuals that are at genetic distance smaller than the threshold are in the same species. Manceau and Lambert (2019) showed that there is a unique finest species partition satisfying the two conditions above, and gave a simple algorithm to find it.

### 1.4 Archetypal models of speciation

From now on, we focus on archetypal models of speciation that rely only on the following three basic assumptions:

1. Speciation is driven by reproductive isolation between individuals or populations, itself resulting from excessive genetic dissimilarity; see Box 2. Typically, individuals are endowed with a partially heritable genotype that can undergo evolutionary changes over time. Genetic differentiation between populations, also called population differentiation, builds up as spatially segregating mutations fix in these populations, due to genetic drift, founder effects or divergent adaptation. Dissimilarity can then be quantified as a measure of the distance between genotypes, and this distance can be used to cluster individuals into species (see Box 1).
2. Genetic dissimilarity is the result of a balance between two processes: spontaneous differentiation driven by mutation as in the previous item, and homogenizing processes such as reproduction and migration. For example, during the time when partially isolated populations are in the “grey zone” (no longer a clear species and not yet two reproductively isolated species, De Queiroz, 2007; Roux et al., 2016), old alleles can spread back and replace new alleles, resulting in the failure of these ephemeral populations to speciate (speciation collapse, whereby incipient differentiation is reset to 0 by the effect of mixing gene pools). Within this framework, reproductive isolation can occur when differentiation predominates over homogenization or when historical contingency factors disrupt the equilibrium between these two antagonistic processes.
3. Dissimilarity can feed back on homogenizing processes, by inhibiting them (e.g., outbreeding depression) or promoting them (e.g., disassortative mating). For example, the ability to interbreed may decrease continuously as a function of genetic dissimilarity, or it may disappear abruptly when dissimilarity exceeds a certain threshold, e.g., representing genetic incompatibilities (Corbett-Detig et al., 2013; Coyne and Orr, 1989; Matute and Cooper, 2021) (see Box 2).

We categorize archetypal models into three classes: **(1) clonal evolution models** study how lineages spontaneously diverge from their ancestor with time, **(2) models of genetic isolation** track the evolution of genetic compatibility between connected populations, and **(3) models of isolation by distance** study how populations freely moving in space become differentiated.

In all cases, differentiation occurs spontaneously as a consequence of mutation but, as we will see, what distinguishes these three classes is the main homogenizing mechanism: stochastic births and deaths (ecological drift) in clonal evolution models, migration between local populations (gene flow) in models of genetic isolation, dispersal and range expansion (spatial drift) in models of isolation by distance.

As explained in the previous section, despite their simplicity archetypal models can model a wide variety of mechanisms (not only neutral) and make specific predictions on a range of macroecological and macroevolutionary observables that can be confronted to real data, for example:

- Interfertility relationships (“who can interbreed with whom”, see Box 1);
- Measures of genomic diversity (see Box 2): genetic diversity within/between species, distribution of genetic differentiation along the genome;
- Measures of species diversity (see Box 3): species richness, species abundance distribution (SAD) – and, in the case of spatial models: range size distribution and spatial distribution of species (species-area relationships (SAR), alpha, beta and gamma diversity);
- Phylogenetic patterns (see Box 4): speciation and extinction rates, phylogenetic balance, lineage-through-time plots, phylogenetic diversity, shape and coupling of gene trees and species trees.

In the next section, we review the main three classes of archetypal models of speciation defined above and their macroscopic predictions. In the last section, we propose some promising avenues of research, conceptual questions to be solved and new models to address them.

#### Box 2

##### The genetic architecture of reproductive isolation.

In models considering the genetic basis of reproductive isolation (RI), individuals are endowed with a haploid or diploid genome with *L* loci carrying alleles taking values in a discrete (usually finite) set. These models mainly fall into two categories.

- **RI as a by-product of genetic distance.** We think of the genetic distance *d* between two genomes *g* and *g^′^* as

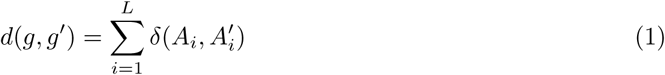

where *A_i_* (resp. *A^′^_i_*) is the allele carried at locus *i* by genome *g* (resp. *g^′^*), and *δ* is a distance in the allele space. Here, the idea is that excessive genetic distance alters the frequency of mating events (premating isolation) or their outcome (postmating isolation), because the accumulation of differences on a locus-by-locus basis increases dissimilarity between phenotypes that can directly (e.g., shape of genitalia) or indirectly (e.g., phenology) be related to reproduction. Nei et al. (1983) used this framework with *L* = 1 or *L* = 2 and a stepwise mutation model, forbidding reproduction when *δ*(*A*_1_*, A^′^*_1_) or *δ*(*A*_2_*, A^′^*_2_) is strictly larger than 1, where *δ*(*A, A^′^*) is the number of mutations needed to go from *A* to *A^′^*. Higgs and Derrida (1991, 1992) introduced a model with two alleles at each locus, forbidding reproduction when *d > d*_min_, with *δ*(*A, A^′^*) = 1*/*2*L* if *A* = *A^′^* (and 0 otherwise).

Due to local adaptation to different environments, genetic differences inevitably arise in key ecological loci which thus effectively act as barriers against gene flow. The “genic view of speciation” (Wu, 2001) predicts that regions of the genome flanking these loci also undergo reduced gene flow, due to effectively reduced recombination close to these loci caused by outbreeding depression, a phenomenon known as divergence hitchhiking (Via and West, 2008; Via, 2009). These regions are called genomic islands of differentiation or genomic islands of speciation (Seehausen et al., 2014; Wolf and Ellegren, 2017).

**RI due to genetic incompatibilities.** Genetic incompatibilities are negative epistatic interactions between alleles at *different* loci affecting the success of mating events, altering for example zygote formation, zygote viability (intrinsic isolation) or hybrid fitness (extrinsic isolation).

The starting point of this approach is the so-called **Dobzhansky-Muller incompatibility** (DMI) (Dobzhansky, 1937; Muller, 1942) involving two loci. The ancestral genotype is *AABB*. In one subpopulation, a mutation is fixed at the first locus (*aaBB*). In another subpopulation, a mutation is fixed at the second locus (*AAbb*). Hybridization would lead to the co-occurrence of incompatible alleles *a* and *b* in the inviable genotype *AaBb*.

Several examples of DMI have been observed in nature. For example, in some marine invertebrates, the two loci correspond to a sperm protein and an egg protein (Vacquier and Swanson, 2011). In the *Xiphophorus* family, they correspond to an oncogene and its repressor: hybrids lack the repressor and develop melanomas due to an excess of melanocyte proliferation (Gordon, 1931; Coyne and Orr, 1989; Patton et al., 2010).

In the case of *L* loci, the most frequent way to model negative epistatic interactions is by a system of *^L^* independent locks associated to all pairs of loci supported by two different-sex gametes. Then the reproductive compatibility *c* between two haploid gametes *g* and *g^′^* can be expressed as

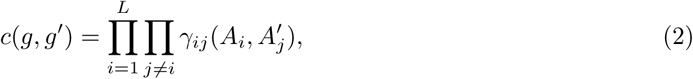

where *γ_ij_* (*A, A^′^*) is the probability that the potential reproductive barrier associated to loci *i* and *j* is inactive when the first gamete carries allele *A* at locus *i* and the second gamete carries allele *A^′^* at locus *j*. Note the contrast of (2) with (1). A common assumption introduced by Orr (Orr, 1995; Orr and Orr, 1996; Orr, 1996) is that all *γ_ij_* (*A, A^′^*) are equal to 1 if *A* and *A^′^* are ancestral alleles, and to some *γ <* 1 otherwise. Then the compatibility equals

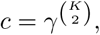

where *K* is the number of mutated loci in at least one of the two gametes, leading to the so-called “snowball effect”: if the number of substitutions increases linearly with time, the number of incompatibilities – and therefore the probability that two individuals cannot interbreed – increases quadratically with time.

The aforementioned models can be understood under the metaphor of “holey” adaptive landscapes suggested by Gavrilets (1997). In this type of fitness landscape, two reproductively incompatible genotypes are connected by a chain of intermediary equally fit genotypes, forming a “ridge”. This makes it possible for two reproductively isolated species to emerge without having to cross a fitness valley. This idea can be extended to higher dimensional genotype spaces (Gavrilets and Gravner, 1997), and more complex models have also been proposed; see Gavrilets (2014) for a review.

#### Box 3

##### Macroecological metrics.

Microscopic models of speciation can offer a better understanding of the mechanisms that underlie commonly observed patterns in macroecology, measured by the following metrics.

**The Species Abundance Distribution (SAD)** in a given community gives the number of species with abundance *n*, for all *n*. A remarkable fact is that most communities’ SAD display a “hollow” shape (Williams et al., 1964; Magurran, 2003), where a handful of species are abundant and most other species are rare (Hubbell, 2001), see the following plot of data taken from Fisher et al. (1943a), which represents species abundance distribution of butterflies collected in Malaya.

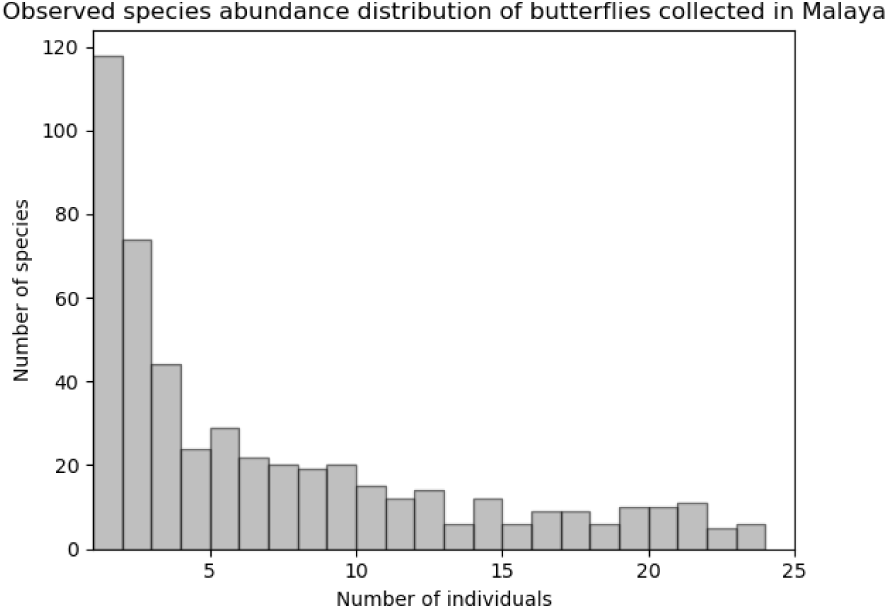

Multiple parametric models have been proposed to fit this hollow curve, such as Fisher’s logseries (Fisher et al., 1943a) or Preston’s lognormal distribution (Preston, 1948), and many other authors have tried to explain the empirical shape of SADs, either via purely statistical or by mechanistic arguments. See e.g. McGill et al. (2007) for an account of the theories.

**The Species-Area Relationship (SAR)** gives the number *S* of species expected to be observed in a geographical zone with area *A*. The most common SAR is called the Arrhenius relationship (Arrhenius, 1921; Preston, 1948) and posits that *S* is proportional to a power of *A*:

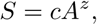

where *c, z* are constants. This relationship is not supposed to hold at all spatial scales. Rather, it is generally believed that SARs are triphasic (Hubbell, 2001; Rosindell and Cornell, 2007) and follow an inverted S-shape when plotted on a log-log scale, displaying a linear intermediate phase with slope *z*. Many other models of SAR exist, see for example Tjørve (2003) for a review.

**Beta diversity** is one of Whittaker’s (Whittaker, 1960) indices of biodiversity, along with alpha diversity which represents diversity at a small (local) scale, and gamma diversity, which represents diversity on a larger (regional) scale. Beta diversity represents the variation of diversity among these different scales, linking local and global species diversity. It is a concept that is quantified in several ways in the literature, which do not always account for exactly the same phenomena, see (Tuomisto, 2010). Whittaker (1960) proposed for example to measure this index by defining it as the ratio between gamma and alpha diversity. Chave and Leigh Jr (2002); Zillio et al. (2005); O’Dwyer and Green (2010) quantify it using the probability that two individuals randomly sampled at a distance *r* belong to the same species.

**The Range Size Distribution** gives the proportion of observed species with a given range size. Speciation rate is thought to be one of the major determinants of range size. However, the relation between range size and speciation rate shows conflicting evidence (Gaston, 1996). Note that under some specific additional assumptions, SAR can be deduced from range size distribution, for example when ranges are approximated as disks and their centers form a Poisson point process in the plane, see Allen and White (2003).

#### Box 4

##### The shape of phylogenetic trees.

Species phylogenies carry some information on the diversification process that has generated them. This information can be quantified by comparing phylogenies or by summarizing them by well-chosen descriptive statistics.

In order to compare phylogenies, one can use a distance function on the space of trees, such as the Robinson–Foulds distance (Robinson and Foulds, 1981) or one of the many available alternatives (see e.g. Kuhner and Yamato 2015). However, in order to uncover general patterns – or simply to focus on specific aspects of phylogenies – it is often convenient to use summary statistics, also referred to as *shape statistics* in that context.

**Balance indices** are one of the most important classes of such shape statistics. Their goal is quantify the intuitive idea that some trees are more “balanced” or “have more symmetries” than others. For instance, in the example below, the complete binary tree on the right conforms more to our idea of what it means to be “balanced” than the caterpillar tree on the left.

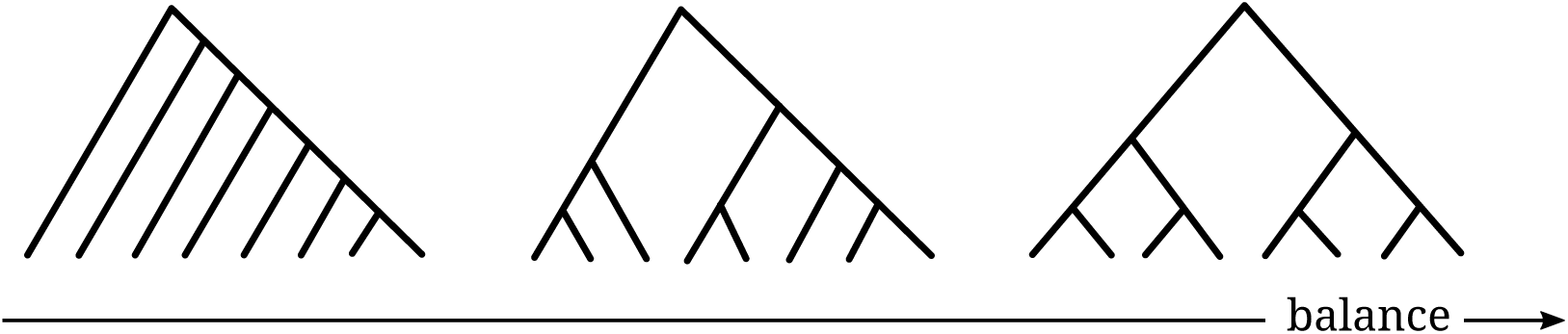

The best known balance indices are perhaps the Colless index (Colless, 1982) and the Sackin index (Shao and Sokal, 1990); but there are about 20 balance indices whose mathematical properties and usefulness in applications have been studied extensively over the past decades, see Fischer et al. (2021 preprint) for a recent survey. In addition to these indices, an alternative approach to quantify the balance of a phylogeny is to fit a model whose parameter is expected to correlate with some notion of balance. Two prominent examples of this are the parameter *α* of Ford’s model (Ford, 2006) and the parameter *β* of Aldous’s *β*-splitting model (Aldous, 1996).

One of the uses of balance indices is to test whether new species appear more frequently in some clades than in others, without any information about the history of unsampled clades or extinct ones. For example, it is known that trees generated by constant-rate birth-death processes are significantly balanced (*β* = 0). Remarkably, most phylogenies encountered in practice turn out to have a comparable degree of balance: significantly lower than that of birth-death trees but significantly higher than that of uniform (“proportional to distinguishable arrangements”) trees (*β* = −3*/*2), often close to a *β*-splitting model with *β* = −1 (Blum and François, 2006). This is an example of a macroevolutionary pattern that has yet to be fully explained.

**The *γ* index and LTT plots** belong to another important class of shape statistics that aim to measure “how early *vs* late” most speciation events occurred (or, more generally when branch lengths do not correspond to physical time, how “close to the root” the nodes of the tree are). For instance, in the time-embedded phylogenies below, speciations tend to occur earlier in the phylogeny on the left than in the phylogeny on the right.

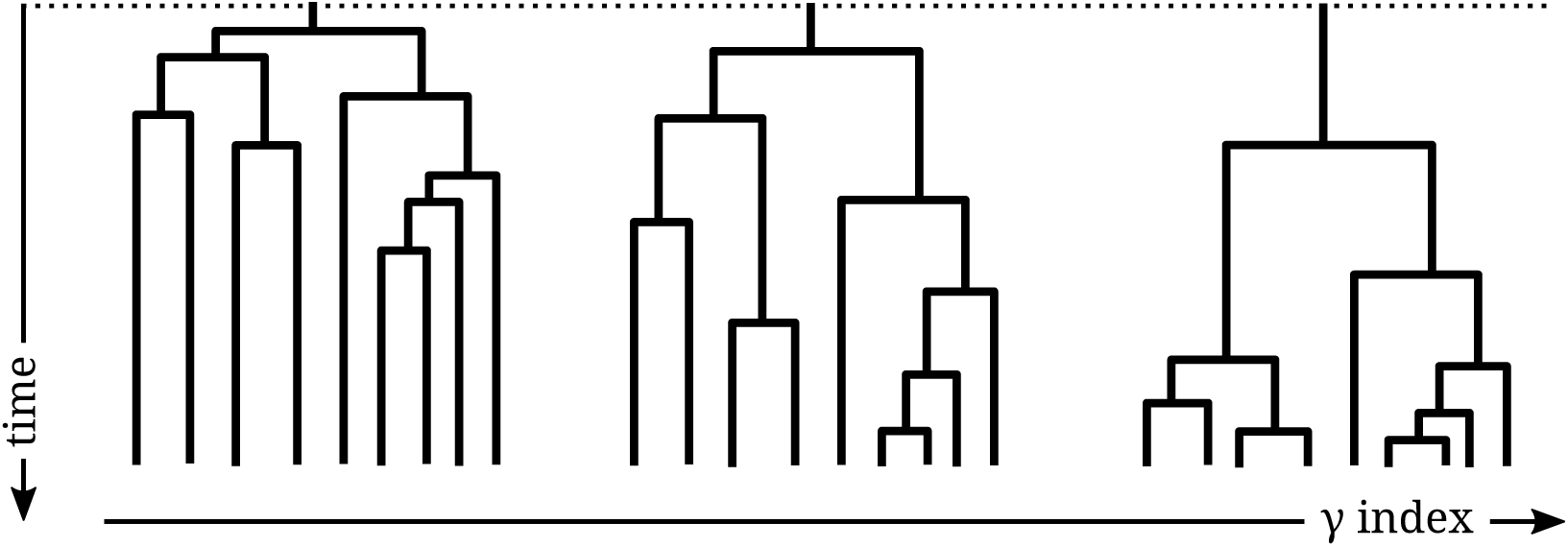

The best known statistic to capture this idea is the *γ* index introduced by Pybus and Harvey (2000); but, to get a more complete picture, lineage-through-time (LTT) plots are frequently used (Harvey et al., 1994). Unlike balance indices, which depend only on tree topology, the *γ* index and LTT plots use branch lengths. Moreover, whereas balance indices quantify how uniformly speciation events are distributed “horizontally” in the tree, the *γ* index and LTT plots provide information about their “vertical” distribution in the tree; they can therefore be used to test, e.g, whether speciation rates are time-homogeneous. In particular, a negative value of *γ* is often observed in practice and can be interpreted as a diversification slowdown (Moen and Morlon, 2014). This is yet another example of a macroevolutionary pattern that is not fully understood.

• **The phylogenetic diversity index (PD index)** aims to measure biodiversity as the evolutionary heritage of a sample of species using their phylogeny. There are many other options to do that (see e.g. Schweiger et al., 2008), but the PD index introduced by Faith (1992) is the most commonly used. It is simply the total branch length of the tree. Besides its use in phylogenetics, the PD index has been used in conservation, as it can help us understand, e.g, how much biodiversity is lost as a result of species extinction (Mooers et al., 2012; Lambert and Steel, 2013).

## 2 A review of archetypal models of speciation

### 2.1 Clonal evolution models

We borrow the terminology of cancer research and immune response initiation, where “clonal evolution” refers to somatic cell lineages that diverge by accumulating mutations. **Clonal evolution models** assume that differentiation between individuals and populations can only increase as genomes mutate in parallel. In turn, speciation can only be slowed down by birth and death events purging this diversity.

In this context, the simplest model of speciation is the *point mutation model* of Hubbell’s *unified neutral theory of biodiversity and biogeography* (UNTB). Hubbell introduces the notion of *ecological drift* (demographic stochasticity and neutral competition between individuals across all species, see Box 5 for a precise modeling assumption). His individual-based model exactly mirrors fixed-population-size neutral models of population genetics, i.e. Moran or Wright–Fisher models with mutation, under the infinite-allele assumption (see Section 1.3 and Figure 2). This model reconciles Fisher’s logseries (Fisher et al., 1943b) and Preston’s lognormal distributions (Preston, 1948) for species abundances (see Box 3), as it predicts a SAD that approaches one or the other distribution as the three parameters of the model vary.

The neutrality assumption underlying Hubbell’s theory has raised strong criticism: for instance Clark and McLachlan (2003) use fossil data to argue in favor of (non-neutral) stabilizing mechanisms in forest ecology; Dornelas et al. (2006) argue that environmental stochasticity drives diversity patterns in coral reefs – see also other references in Etienne et al. (2007). Furthermore, among researchers who do not reject the neutrality assumption, Hubbell’s theory has been criticized for the lack of detail in modeling speciation – notably the fact that in UNTB, speciation is instantaneous and occurs at a rate exactly linear in species abundance. Etienne and his co-authors have proposed more flexible versions of Hubbell’s initial model: Etienne et al. (2007) proposed a modification of the theory where the speciation rate is constant across species, as in the birth-death models introduced in Section 1.1, and Etienne and Haegeman (2011) introduced a model of speciation by *random fission* – a first attempt at modeling allopatric speciation, see Box 5. Interestingly, both models failed at improving the fit to SAD data previously provided by the point mutation model. Assumptions on the mode of speciation thus have an important impact on species abundance distributions – see Kopp (2010) for a more complete review on this subject.

With the increasing number of phylogenetic trees being reconstructed through molecular methods, Jabot and Chave (2009) proposed to take phylogenies into account when fitting data to Hubbell’s neutral model. They were able to implement this using an approximate Bayesian computation (ABC) method thanks to the (relatively computation-efficient) representation of genealogies in UNTB as coalescent processes – see Wakeley (2004) for an introduction to coalescent theory. Because the so-called fundamental biodiversity number *θ* (the rescaled speciation rate, see Box 5) impacts phylogenetic balance (see Box 4), their method improved the estimation of the UNTB parameters compared to existing methods. However, it has been argued that this model makes unrealistic predictions about species lifetimes, speciation rates and number of rare species by Rosindell et al. (2010), who introduced a model where speciation is not instantaneous. This model, known as the *protracted model of speciation*, is a multistage model where species are formed of a dynamic swarm of populations that are able to evolve into so-called “good” species after surviving long enough (see Box 6). The model yields realistic predictions for quantities related to species lifetimes, as well as for the SAD. Its predictions on phylogenies have been studied by Etienne and Rosindell (2012); Etienne et al. (2014) and Lambert et al. (2015), who proposed it as an explanation for the diversification slowdown near present-time observed in real phylogenies. In particular, Etienne et al. (2014) showed that the protracted birth-death diversification model correctly estimates the waiting time to speciation from phylogenies.

An advantage of the previous models is that the genealogy of individuals and the clustering of the population into species (see Section 1.3) are constructed jointly. However, the resulting species partitions are not monophyletic in general (Manceau et al., 2015; Manceau and Lambert, 2019), i.e. a species cannot always be defined as the group consisting of all descendants from a single ancestor. In order to circumvent this issue, Manceau et al. (2015) proposed another definition of species in this context, yielding a coarser clustering into species, with a method leading to what they called a model of *speciation by genetic differentiation*. Given a set of point mutations distributed across a genealogy, species are defined as the smallest monophyletic groups of individuals such that any pair of individuals carrying the same genotype are always in the same species. With this criterion, speciation takes time, as mutations arising in a large species will generally not generate a new species instantaneously – a property that makes a bridge with protracted speciation models. The authors found that this model yields realistic phylogenies – i.e. with *β* and *γ* statistics (see Box 4 and references therein) close to those observed in real data, provided that communities are assumed to be expanding – that is, when the constant-population-size (also called zero-sum) assumption of Hubbell’s UNTB is relaxed.

#### Box 5

##### The Unified Neutral Theory of Biodiversity.

The *Unified Neutral Theory of Biodiversity* (UNTB) is a conceptual framework introduced by Hubbell (2001) that regroups several closely related models. In its simplest form, it describes the diversity and abundances of species on two scales: in a *local community* of size *J*; and in a much larger *metacommunity* of size *J_M_*.

The metacommunity dynamics are governed by two phenomena:

- **Ecological drift.** Each individual dies at constant rate and is then replaced by the offspring of another individual sampled uniformly at random.
- **Speciation by point mutation.** Each individual mutates and starts a new species at rate *µ*.

The equilibrium distribution for the metacommunity displays a logseries SAD, where the *fundamental biodiversity number θ* = 2*µJ_M_* takes the role of Fisher’s *α* parameter. This equilibrium state is known as *Ewens’ sampling formula* in population genetics.

Assuming that the mutation rate is small – i.e. *µ* = *θ/*(2*J_M_*) 1 – and that the local community is much smaller than the metacommunity (*J J_M_*), it becomes possible to neglect speciation in the local community. Its diversity then comes through immigration from the metacommunity, which acts as an external buffer. More specifically, the local community dynamics are governed by the following events:

- **Ecological drift and immigration.** Each individual dies at constant rate and is then replaced by the offspring of another individual. With probability *m*, this individual is sampled uniformly in the metacommunity; with probability 1 − *m*, it is sampled uniformly at random in the local community.

The equilibrium species abundance distribution in the local community is called the zero-sum multinomial distribution. The lower the immigration parameter *m*, the more isolated the local community and the fewer rare species (i.e. species with few individuals) at equilibrium, which results in a lognormal-like SAD.

In this model, when new species appear they only consist of a single individual. Hubbell (2001, 2003) and Hubbell and Lake (2003) proposed alternative modes of speciation:

- **Speciation by random fission.** At each speciation event, a species of size *N* is divided into two new species: one of size *K*, where *K* is a uniform random variable between 1 and *N*; and the other of size *N − K*. This mode could represent a random geographic barrier that splits the species into two.
- **Peripatric speciation.** An intermediate mode between point mutation and random fission, where new species start with a small number of individuals.

#### Box 6

##### The protracted model of speciation.

Rosindell et al. (2010) criticized previous models of neutral speciation for failing to take into account the fact that speciation takes time. Their model of *protracted speciation* addresses this by considering a parameter *τ* that corresponds to the time it takes for a so-called *incipient species* – i.e. a subpopulation of a species that has started to differentiate – to turn into a species in its own right (a so-called *good species*). Behind this transition period are hidden complex biological processes that the authors do not model explicitly (see Section 3.1). In particular, this allows the authors to take into account the time it takes to form new species, without explicitly modeling the genetic mechanisms explaining this gradual speciation and without bookkeeping all pairwise differences between genomes.

The basic model is an extension of Hubbell’s UNTB model described in Box 5. The local community dynamics are unchanged, but the metacommunity dynamics are replaced by the following process:

- **Ecological drift.** Each individual dies at constant rate 1 and is immediately replaced with a offspring of another individual selected uniformly at random.
- **Differentiation by point mutation.** Each individual mutates at rate *µ*: it is replaced with an individual forming a new incipient species.
- **Protracted speciation.** After a transition time *τ*, an incipient species becomes a good species.

In this model, *µ* is to be interpreted as a “speciation initiation” rate, and the true speciation rate is *µ*_eff_ = *µ/*(1 + *τ*). In the metacommunity, the model predicts an expected number of species with abundance *n* of

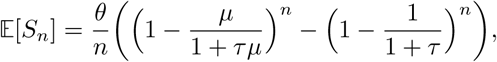

where *θ* = *µJ_M_ /*(1 + *µ*) – this yields a logseries-like SAD for small values of *τ* and a lognormal-like SAD for large values of *τ*.

**Protracted birth-death models.** In the setting of growing populations, another approach to protracted speciation – proposed by Etienne and Rosindell (2012) and further studied by Etienne et al. (2014) and Lambert et al. (2015) – makes the time to speciation a random time: a species gives birth to an incipient species at rate *λ*_1_, which turns into a good species at rate *λ*_2_. The authors show that these models yield satisfying predictions about the shape of phylogenetic trees, in particular they can explain the diversification slowdown near the present (Etienne and Rosindell, 2012), and they accurately predict the waiting time to speciation (Etienne et al., 2014).

### 2.2 Models of genetic isolation

In **models of genetic isolation**, we explicitly consider whether two individuals or populations are reproductively compatible. Doing so allows us to define species directly from interfertility relationships, e.g. as the connected components of the graph whose vertices are populations and edges represent interfertility (see Box 1). Bienvenu et al. (2019) considered such a representation explicitly: a random graph in which edges vanish after some random time, mimicking the buildup of reproductive isolation; and evolving under extinction-recolonization dynamics, where each newborn population inherits its neighbors from its mother. They make accurate predictions on the vertex degree and the number of species, but further work is needed to provide joint predictions for phylogenies and species abundances (see Boxes 3 and 4).

Most other models build on clonal evolution models and on additional assumptions on the genetic basis of reproductive isolation. These models generally use an explicit representation of genomes, with varying degrees of faithfulness (number of genes, physical linkage, etc.).

In some, but not all, in addition to mutations differentiating gene pools, gene flow (sexual reproduction, migration between demes) can tend to homogenize them. This gene flow can be maintained as long as the dissimilarity between individuals or populations remains low enough. In the most complex models, increasing dissimilarity due to mutations is assumed to in turn reduce gene flow, potentially leading to reproductive isolation. One could also consider cases where dissimilarity enhances gene flow (e.g., disassortative mating, self-incompatibility) but these models are not studied here.

We now review models where individuals are endowed with an explicit genotype and two individuals are declared reproductively isolated based on the dissimilarity between their respective genotypes. Recall from Section 1.3 how to define species under this assumption. As specified in Box 2, we can distinguish between two subcategories of models depending on assumptions on the genetic basis underlying reproductive isolation: reproductive isolation as a by-product of genetic distance or reproductive isolation due to genetic incompatibilities.

#### 2.2.1 Reproductive isolation as a by-product of genetic distance

The main example of an archetypal model of distance-based speciation with gene flow is the *Higgs– Derrida model* (HDM) of speciation (see Box 7). Higgs and Derrida (1992) considered a panmictic population of fixed size, with a large number of loci undergoing mutation and recombination, and proved that speciation could occur without selection nor geographic structure. Here, reproductive isolation occurs when the genetic distance exceeds a threshold, see Section 1.3 and Boxes 1, 2 and 7. It is natural to embed such a model (Higgs and Derrida, 1991; Nei et al., 1983) in a metapopulation composed of demes connected by migration. For example, Manzo and Peliti (1994) have extended the HDM to a model where individuals live on islands connected by rare migration and showed that speciation is more likely in their model than in the original sympatric HDM.

The assumption of rare mutations and rare migrations is commonly used to overcome difficulties caused by the potentially large number of coexisting genotypes, as it makes it possible to separate timescales and to neglect variations within populations. The resulting drastic reduction of dimension allows for exact analytic approaches, leading to fruitful predictions. For example, Gavrilets (2000) proposed a deterministic approximation of the dynamics of the genetic distance between the mainland and a peripheral population subject to immigration, assuming that the population is at all times monomorphic with respect to the loci controlling incompatibilities. This approach makes it possible to estimate the speciation time, defined as the time when the genetic distance reaches a predefined threshold, depending on model parameters (migration and mutation rates, threshold value).

A similar methodology has been adopted by Yamaguchi and Iwasa (2013), who introduced a stochastic extension of Gavrilets’s deterministic model that accounts for the randomness of migrations and mutations. They were able to find analytical results on the waiting time to speciation, depending on the initial genetic distance between populations and on the threshold for genetic isolation, and compare it with individual-based simulations. Notably, even in cases where the deterministic approximation suggests that the genetic distance will stabilize below the speciation threshold – so that migration is strong enough to prevent speciation – the occurrence of a rare sequence of migration events could potentially drive the population towards genetic incompatibility and thus decrease the waiting time to speciation.

Most metapopulation models focus exclusively on a small number (2 or 3) of islands. Miró Pina and Schertzer (2019) introduced a generalization of Yamaguchi and Iwasa (2013) to a general metapopulation model, to understand how more intricate geographical constraints might impact the aforementioned predictions. One conclusion of this approach is that the steady-state structure of the metapopulation into species is heavily influenced by the quantity of potential migration pathways connecting any two islands, and that a greater number of clusters or geographical bottlenecks are more likely to facilitate speciation events.

##### Box 7

###### The Higgs–Derrida model of speciation.

Higgs and Derrida (1991, 1992) introduced a model where a single population of size *M* is studied, subject to sexual reproduction and mutation. Each individual of this population bears a genome *g* of size *L*, which consists of a family of “spins” (*A^g^_1_, A^g^_2_, …, A^g^_L_*), with *A^g^_i_* ∈ {−1, 1} for all *i*. Generations are non-overlapping and the population size is constant, similarly to the Wright-Fisher model.

This model takes into account the genetic distance between each pair of genomes *g* and *g^′^* using the notion of *overlap q^g,g′^* = 1 − 2*d*(*g, g^′^*), where *d*(*g, g^′^*) is the distance between *g* and *g^′^* defined as:

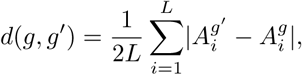

which equals ^1^ (in mean, and exactly when *L* is brought to infinity) when the two genomes are independent, and 0 when the two genomes are equal.

The genomes of a new generation of individuals are generated using a few key ingredients:

- **Sexual reproduction:** At each generation, each individual, independently, chooses two parents whose genomes are at a distance lower than a fixed threshold 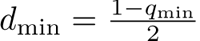.
- **Free recombination and mutation:** At each locus independently, the individual inherits the allele of one of its two parents with probability ^1^. This allele then mutates with probability *µ*. Species are then defined by a threshold clustering algorithm with threshold *d*_min_ (see Section 1.3). Higgs and Derrida (1992) numerically simulated the evolution of the overlaps when *L* is large, starting from a homogeneous population. In particular, they were able to make estimates on:
- **Relation between species abundance, mutation rate, and speciation:** If 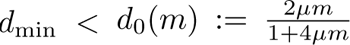, then any species of size *m* is going to split into two species. Therefore the condition to get at least one speciation event is *d*_min_ *< d*_0_(*M*).
- **Inter/intra-specific diversity:** For each pair of species *A* and *B* of respective sizes *m_A_, m_B_*, one can define 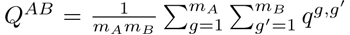 the mean pairwise overlap between *A* and *B*.

Taking *A* = *B* gives an idea of the intraspecific diversity of *A*, whereas *A* ≠ *B* gives an idea of the interspecific diversity between the two species (diversity decreases with overlap). Another indicator of intraspecific diversity is the function *σ_A_* which records the mean number of pairs of individuals which are not able to interbreed inside species *A*. Simulations indicate that *σ_A_* stays close to 0 most of the time, except during short speciation events. This seems to validate the choice made for the species definition in this model.

#### 2.2.2 Reproductive isolation due to genetic incompatibilities

In the classical representation of the two-locus Dobzhansky–Muller incompatibilities (DMI), two allopatric populations start out with the same genetic composition *AABB* (see Box 2). Most models focusing on this setting study the waiting time before the appearance of each mutant *a* and *b* in its background and its fixation (to *aaBB* and *AAbb*, respectively), or the conditions under which total isolation is eventually completed in spite of gene flow (Bank et al., 2012; Blanckaert and Hermisson, 2018; Gavrilets, 1997; Gavrilets and Gravner, 1997).

Most generalizations of this model to multiple (partially) interacting loci also focus on two populations in allopatry and the time to speciation in the absence of gene flow. Orr and Turelli (2001) extended the seminal paper by Orr (1995) (which introduces the so called “snowball effect”, see Box 2) by studying the time-evolution of the number of DMI when the number of substitutions between the two populations is itself random and incompatibilities’ effects differ, defining speciation time as the time when this number reaches some threshold. The case of a finite number of loci and potentially decreasing genetic distance is studied by Palmer and Feldman (2009).

Other studies look at the influence of different incompatibility scenarios, and the likelihood of the snowball effect (Cutter, 2012; Livingstone et al., 2012; Gourbiere and Mallet, 2010; Fraïsse et al., 2014; Kondrashov, 2003). For example, Livingstone et al. (2012) studied the probability of speciation in the BDM model as a function of the complex protein-protein interaction network and the probability of deleterious interactions, thus taking into account the fact that some pairs of proteins do not interact. They showed in particular that empirical networks produce lower rates of speciation than the denser complete networks.

The DMI framework relies on a very simple fitness landscape, called holey landscape (see Box 2), where crosses from peaks (double heterozygotes) fall into the fitness valley (are inviable). Several authors have focused on a specific instance of this situation caused by chromosomal variants such as inversions or translocations starting with Wright (1941) and followed in particular by Lande (1979); Coyne et al. (2000); Rieseberg (2001); Kirkpatrick and Barton (2006).

It is not known what can happen in a setting where incompatibilities themselves are dynamic, so that new species can constantly arise. Marin et al. (2020) made a first attempt at modeling this, by assuming that introgression of a gene into a population can occur with a probability that increases with the number of co-adapted alleles carried by all other loci in the receiver genome. The main assumption is that each novel allele arising by mutation is co-adapted with all alleles of the background where it arose (or is purged by purifying selection) – but with no alien allele. This embodies the idea that a novel allele needs to be “tested” against the genome harboring it.

The model enforcing this assumption, called *gene-based diversification model* (GBD), makes some specific predictions on the joint evolution of gene and species lineages, acknowledging that gene trees can greatly differ from each other (and, therefore, from the species tree).

The assumptions of this model can be justified when migration is dominated by unlinked genes. In this case, papers such as Barton and Bengtsson (1986) and Westram et al. (2022) show that even in models with explicit hybrid fitness depression, the effective migration rate can be computed (as the product of mean fitnesses of successive backcross generations). This suggests that the range of validity of archetypal models like GBD goes well beyond simple, neutral mechanisms.

### 2.3 Models of isolation by distance

A key element missing from most of the previous models is the explicit geographical context of speciation. Spatially explicit models are of particular interest for predicting and understanding empirical observations such as species-area relationships and range size distributions (see Box 3). **Models of isolation by distance** consider spatially embedded populations and seek to study the build-up of species boundaries in space. We discard from this category static models of metapopulations composed of a few discrete demes connected by migration. In contrast, we assume here that individuals are positioned in large, regular grids or in continuous space where distance can be physically measured. In these models, species emerge as a result of isolation by distance in addition to possible other mechanisms such as genetic differentiation.

A first way to construct a spatially explicit model of speciation is to extend an existing nonspatial model, such as those introduced by the UNTB (Box 5). This can be done for example by introducing a dispersal kernel (see Box 8), as in Durrett and Levin (1992), who used a contact process, i.e. a stochastic nearest-neighbour epidemic process on the lattice – a natural choice for modeling dispersal. In their models, new species are introduced by immigration from outside the system or by speciation events, the latter being modeled by point mutations. Their objective was to make sense of Arrhenius SAR – i.e. of the fact that the number of species is proportional to a power of the area (MacArthur and Wilson, 1967, see also Box 3). The model predicts that the exponent in the SAR depends on the rate of introduction of new species, which can explain the variability observed across different taxa and locations. Subsequently, Chave and Leigh Jr (2002); Zillio et al. (2005) used models of neutral biodiversity with limited dispersal close to the one of Durrett and Levin (1992) to predict species beta diversity (defined there as the ratio between regional and local species diversity) in tropical forests. Rosindell and Cornell (2007) also used a spatially explicit version of Hubbell’s neutral model where they varied the dispersal range and the dispersal kernel. They found that the model always predicted the same power law for the SAR, up to rescaling of the two axes, and showed that it exhibits the triphasic behavior that is empirically observed in data. Using fat-tailed dispersal kernels Rosindell and Cornell (2009) improved the fit for the SAR, finding realistic values for the exponent of the Arrhenius relationship (Box 5). O’Dwyer and Green (2010) found a first analytical prediction of the triphasic behavior of the SAR in a neutral setting, using quantum field theory, along with a prediction of the beta diversity close to the one of Chave and Leigh Jr (2002). Moreover, they were able to link the exponent *z* in the Arrhenius SAR to parameters of their formula for beta diversity.

An alternative way to spatialize speciation is to start with a model of genetic isolation, as in the previous section. Desjardins-Proulx and Gravel (2012a) extended Economo and Keitt (2008)’s model based on the interactions within a set of spatially-organized local populations under ecological drift (see Box 5), and where reproductive isolation is modeled by genetic incompatibility (see Box 2). Their neutral model could not match empirical data of species diversity with realistic mutation rates. Using random geometric networks in a “pseudo-selection” model, Desjardins-Proulx and Gravel (2012b) found that species richness was higher in more connected communities, while speciation was facilitated in more isolated communities. This result is similar to the prediction made by Miró Pina and Schertzer (2019).

Other models are extensions of the HDM (Box 7) in which individuals or populations also have a spatial location and reproduce with their neighbors. Using an approach close to Manzo and Peliti (1994)’s, Gavrilets (1999) analytically studied the evolution of the mean genetic distance within and between subpopulations in a stepping-stone model, under multiple geographic scenarios (isolated populations, populations linked by migration, peripheral population). Gavrilets et al. (1998, 2000b) used individual-based simulations of this model to make predictions on the waiting time and the location of the first speciation event in relation with local population sizes, mutation rates, dispersal ability and the threshold for reproductive isolation. They notably showed that gene flow does not impede speciation, even in the absence of any mechanism favoring divergence like differential adaptation. Gavrilets et al. (2000a) studied a 1D array of demes undergoing extinction and recolonization, where speciation is modeled by clonal evolution. Focusing on the movement of species boundaries, they obtained analytical predictions for <u>the</u> average number of species and the average species range, showing that the former scales like *δ/µ*, *δ* being the extinction-colonization rate and *µ* the mutation rate per deme. Using numerical simulations, they obtained species range distributions that match those observed in empirical data. de Aguiar et al. (2009) and de Aguiar (2017) also considered a spatial version of the HDM to get what they named a “topopatric” model of speciation. They also found that SAR and SAD match observations made for different taxa and regions. In their models, individuals can only mate if their spatial distance and genetic distance are lower than fixed values. This is something that Martins et al. (2013) also did, in the case of ring species.

In these models, reproductive isolation arises when dissimilarity exceeds a fixed threshold. In contrast, Hoelzer et al. (2008) used cellular automata to explicitly model individuals’ genomes on a 2D grid, in a sexually reproducing population with limited dispersal, where the fitness of an offspring is a decreasing function of the dissimilarity of its parents’ genomes. Pigot et al. (2010) modeled how species boundaries move in a more coarse-grained way than Gavrilets et al. (2000a), overlooking the microscopic dynamics of individuals within species. They considered two modes of speciation: vicariance – when a geographic barrier intersects the range of an extant species – and peripatry – when the individuals in the edge of the range move to a new location. Their predictions on phylogenetic tree shapes – which are imbalanced and show a slowdown in the diversification rates match bird phylogenies; but their predictions on the SAR do not match empirical observations.

#### Box 8

##### Spatial models using a dispersal kernel.

Models of isolation by distance frequently use a *dispersal kernel p* which quantifies the probability density *p*(*x, y*) for a parent located at *x* to disperse its offspring at some other point *y*. This parent-offspring displacement can be:

1. restricted to the parent’s neighbors (as in the contact process, see e.g., Durrett and Levin, 1992; Zillio et al., 2005 or section 3.3)
2. or spread over a larger area around parent (e.g., Rosindell and Cornell, 2007; Chave and Leigh Jr, 2002).

In the first case, individuals will typically be found on points of a lattice (e.g., the two-dimensional grid) and for any pair *x*, *y* of points of this lattice, *p*(*x, y*) will be zero when *y* is not a neighbor of *y*. For the neighbors *y* of *x*, when the space is *d*-dimensional, *p*(*x, y*) can for example be taken equal to 1*/*2*d* in order to model isotropic dispersal. In the second case, space can either be continuous or discrete and *p* can take many forms. A common one in continuous space which is isotropic is the *Gaussian kernel*:

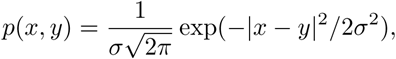

with |*x* − *y*| the Euclidean distance between *x* and *y*. The Gaussian kernel also gives the law of the position of a symmetric random walker after a large number of small steps (Central Limit Theorem). It is remarkable here that gene flow is controlled by only one parameter *σ*, which represents the typical dispersal distance.

## 3 New models and perspectives

In this section, we outline several new archetypal models of speciation, hopefully tractable from a mathematical point of view. Each of these models aims to address a specific question that is only partially addressed by existing models.

The first group of models, which can be seen as refinements of existing protracted speciation models, aim to model **speciation collapse due to secondary contact**, where the genetic difference between two populations can be reset to 0 due to gene flow. They open the black box of “protracted speciation” in a parsimonious way: although gene flow is not modeled explicitly, they allow for incomplete reproductive isolation between populations that have not fully evolved into distinct species yet, and could therefore merge back into one species.

The second group of models aims to address a shortcoming of the threshold models of reproductive isolation reviewed in Section 2.2. Indeed, these models – in which individuals/populations lose all ability to interbreed once their genetic distance exceeds a certain value – do not incorporate the fact that **dissimilarity feeds back on homogenization**: either two populations are similar and can interbreed; or they are dissimilar and cannot interbreed. The model that we introduce to tackle this issue instead uses a continuous relationship between gene flow and dissimilarity.

The third class of models that we introduce are models that aim to make predictions on **spatial patterns of speciation** – in particular, on the relationship between species range and speciation rates. We consider two such models: a purely neutral one (the *freezing voter model*) and a purely adaptationist one (the *Red Queen model*). Despite the simplicity of their formulation, these models, especially the freezing voter model, are challenging to study mathematically.

### 3.1 How does speciation collapse slow down diversification?

One of the innovations of Rosindell et al. (2010)’s protracted speciation model has been to recognize and incorporate the fact that speciation is not instantaneous. However, in this model, the process by which so-called *incipient* species turn into *good* species remains a black box, and is reduced to its duration *τ*, which is set as an exogenous parameter of the model. In a more realistic context, the transition from incipient to good species is the result of a complex interplay between differentiation and homogenization. This process not only unfolds over time, but also carries the potential for failure – resulting in what is referred to as *speciation collapse*.

To gain a comprehensive understanding of this effect, it becomes essential to enhance the initial protracted speciation model with a thorough description of the competition between differentiation and homogenization. This will make it possible to obtain the transition time *τ* of Rosindell et al. (2010) as an emergent property, and at the same time to assess the role of speciation collapse in speciation.

A simple model consists in considering a collection of populations where differentiation is driven by point mutations; homogenization happens as a result of reproductively compatible populations merging; and populations become reproductively isolated past a certain threshold. More specifically, one can for instance consider a set of populations carrying types, where:

- Populations can split or become extinct as in a birth-death process;
- Each population mutates at rate *µ*, taking a new type never seen before (infinite-allele model).
- Each pair of reproductively compatible populations merges at constant rate, where:

**–** Two populations are reproductively compatible if their types are at distance less than *k* in the genealogy of the types (see Section 1.3).

**–** When two populations merge, the type of the resulting population is chosen according to some specific rule. For instance, the merged type could be one of the two parent types chosen at random; or it could be a new type, seen as a child of the two parent types – in which case the genealogy of the types is given by a directed acyclic graph instead of a tree – see Figure 3.

**Figure 3:**
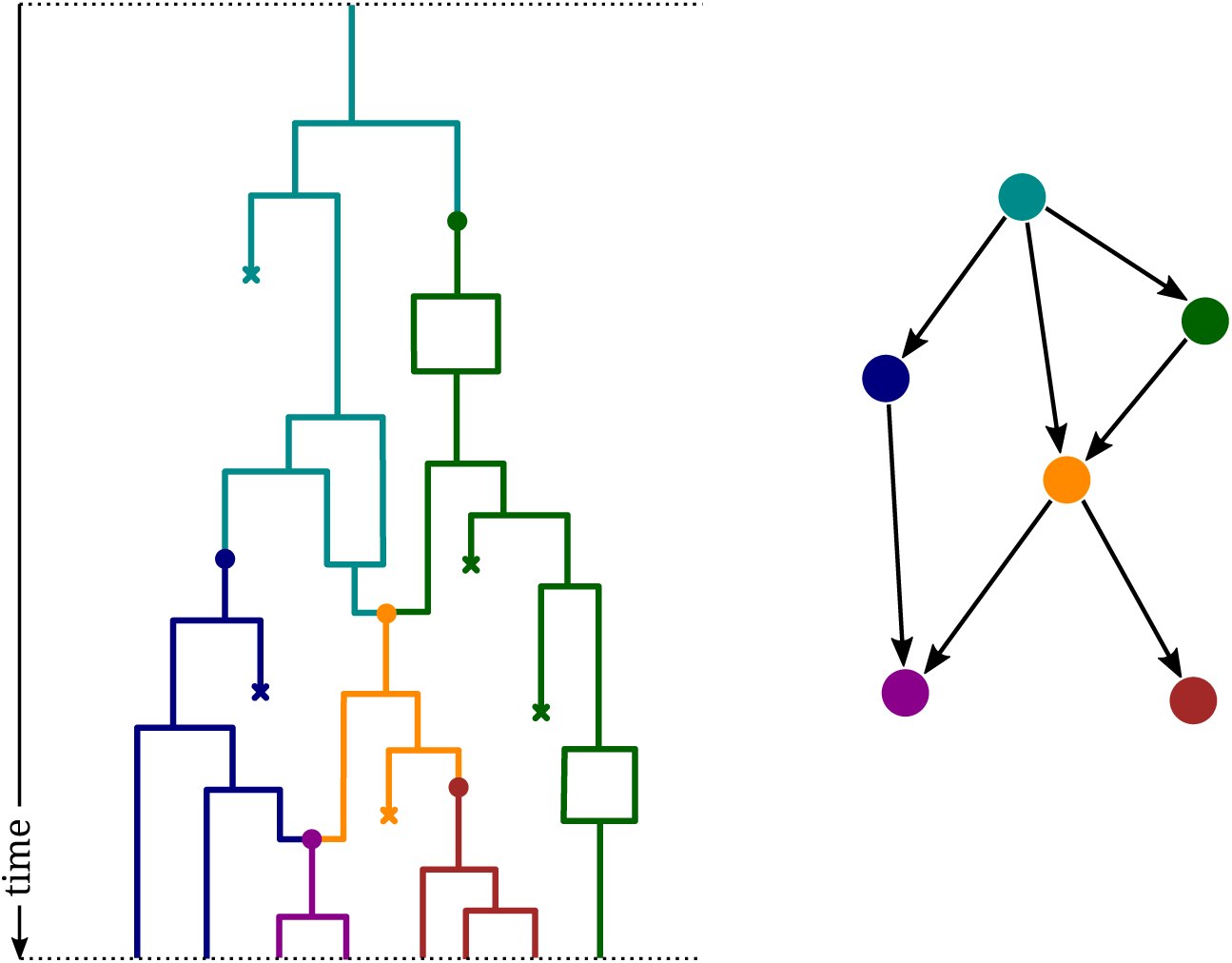
Left, a realization of the process described in the main text. Each vertical line corresponds to the lineage of a population, and each color to a type. Right, the directed acyclic graph encoding the genealogy of the colors. In this example, populations are reproductively compatible if and only if their colors are separated by at most two edges in the genealogy of the colors.

Like protracted speciation models, this model describes the dynamics of a swarm of evolving populations. In protracted speciation models, these populations are purely diverging, and this divergence is artificially described as a two-step process (incipient species turning into good species). Here, not only is there a possibility for homogenization, but divergence is also a more gradual process: there is no need for a notion of incipient species. However, to fall back on the protracted speciation setting, each new type can be interpreted as an incipient species.

The case *k* = 1, where only populations of the same type are reproductively compatible, has been studied in detail in Bienvenu and Duchamps (2024). Even though, like the protracted speciation model of Rosindell et al. (2010), this specific case lacks the crucial ingredient that is homogenization, it provides a helpful guide to see the kind of mathematical results that one can hope to obtain for such models, as well as the challenges that their study poses. In particular, for *k* = 1 differentiation is always sufficient to ensure speciation – in the sense that the network describing the genealogy of the populations has a tree structure on the large scale; moreover, this tree structure is fully understood (namely, it is the continuum random tree of Aldous, 1991). But whether this is the case for every value of *k* is an open and challenging mathematical question.

Another very natural way to model competition between differentiation and homogenization is to assume that populations diverge at a constant rate and that homogenization is therefore a decreasing function of their divergence time. Formally, one can consider a model where, as previously, populations branch and die at a constant rate; and where two populations *i* and *j* merge at rate *f* (*t*_MRCA_(*i, j*)), where *t*_MRCA_(*i, j*) is the time to a most recent common ancestor of *i* and *j*, and *f* is a decreasing function parametrizing the model.

As for the previous model, we say that there is speciation when the network describing the genealogy of the populations has a tree structure on the large scale (i.e. when there are groups of populations that have diverged so much that their descendants coexist without ever merging together). The outcome of the competition between differentiation and homogenization – i.e. whether the homogenization, as parametrized by a given function *f*, is strong enough to prevent speciation – is highly non-trivial.

### 3.2 How does dissimilarity slow down homogenization?

In order to motivate the next model, we recall that in many of the models presented in Section 2.2.1, reproductive isolation was presented as a threshold effect: two individuals are reproductively compatible if their genetic distance is below a predetermined threshold; otherwise they are reproductively incompatible.

In reality, speciation is inherently characterized by a gradual transition, and there is a need to explore the potential outcomes of depicting reproductive isolation as a continuous function of genetic distance. In this model, speciation will not be represented as a sudden, discontinuous process; rather, it will be represented as a gradual fragmentation resulting from the feedback between migration and reproductive isolation. As two populations become more genetically distant, gene flow decreases, consequently leading to an increase in genetic divergence due to the accumulation of new mutations and so on and so forth. Contrary to the threshold model, it is rather unclear whether such a snowball effect will eventually drive two semi-isolated populations into two distinct species.

In order to convince the reader of the potential of such an approach, we revisit the multiscale model alluded to in Section 2.2.1 in more detail. The population is partitioned into demes connected by migration. In this setting, mutations and migrations are rare, so that intra-deme diversity can be ignored. We assume the existence of *L* ≫ 1 speciation loci and define the genetic proximity *p_ij_* between demes *i* and *j* as the fraction of loci where *i* and *j* share the same allele. We assume the existence of a *feedback function h* that encodes the degree of reproductive compatibility as a function of genetic proximity. It is natural to assume that *h* is increasing (i.e. the gene flow between two demes increases with their genetic proximity) and that *h*(0) = 0. Finally,

- In each population and at each locus, new mutations fix independently at rate *µ*;
- An “effective” migration between *i* and *j* occurs at rate *m_ij_ h*(*p_ij_*), where *m_ij_* is the maximal migration rate from *i* to *j*. Upon effective migration, a single allele of the migrant *i* is fixed in population *j*.

The feedback function *h* may be hard to measure in practice, but some of its qualitative features could shed light on potential speciation scenarios.

As an illustration, consider the simple situation of two demes, with genetic proximity *p*(*t*) at time *t*. When *L* ≫ 1 and assuming symmetric migration at rate *m*, the dynamics of *p* are well approximated by a deterministic equation

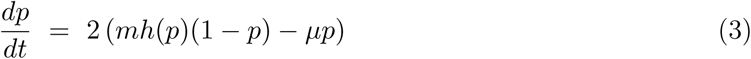

To gain some intuition on the derivation, the first term on the right-hand side is the effect of migration on genetic diversity (*p* increases), whereas the second term is the effect of mutation (*p* decreases). For a general migration graph, the previous approach can be generalized so that the genetic proximities (*p_ij_* (*t*)) are now solution to an explicit, non-linear system of differential equations.

A careful analysis of the limiting deterministic system can enable us to connect the resilience of a species (or conversely, the occurrence of speciation) to the behavior of the feedback function *h* at 0. If *h*^′^(0) *>* 0, the system rebounds from any environmental stress (speciation collapse). If *h*^′^(0) = 0, speciation takes place following sufficiently strong environmental stress.

In summary, the previous model suggests that the feedback function *h* has the potential to yield diverse qualitative predictions on speciation (speciation collapse, ring species). Consequently, it would be intriguing to explore how *h* arises from a more detailed population description. In this refined model, migration is not portrayed as an instantaneous event but rather as a stochastic process, wherein the genetic material of a single migrant gradually disperses over several generations until the fixation of some of its alleles occurs in a focal population. In turn, this invasion of the migrant’s genetic material depends on the specific mechanism of genetic isolation at hand. An interesting question would be to find an approximate relation (if it exists) between a given mechanism of reproductive isolation and the feedback function *h*. Works addressing the effect on effective migration of polygenic divergent adaptation could prove valuable in this perspective (Szép et al., 2021; Sachdeva, 2022). Provided such a program could be achieved, one could then relate speciation patterns to the underlying genetic architectures of Box 2.

### 3.3 How does species range control speciation?

Understanding how species range influences speciation rates is notoriously difficult. Large species ranges can be thought of as favoring speciation, but a large range can be due to a high dispersal ability, which will in turn increase gene flow and reduce population differentiation, thereby inhibiting speciation. Similarly, spatially fragmented species are seemingly more prone to speciation (see for example Smyčka et al., 2023; Ciccheto et al., 2024), but if local populations are too small, they can go extinct before adapting to the local environment. On the empirical side, studies on how speciation is influenced by population size, population structure, species range, evolution of reproductive isolation or population differentiation abound: in gastropods (Wagner and Erwin, 1995), mollusks (Jablonski and Roy, 2003), birds (Harvey et al., 2017; Rabosky and Matute, 2013), lizards (Singhal et al., 2018), desert snakes (Alencar and Quental, 2019), freshwater fish (Yamasaki et al., 2020), drosophila (Rabosky and Matute, 2013) or *in silico* (Birand et al., 2012; Maya-Lastra and Eaton, 2021 preprint; Pigot et al., 2010). Overall, empirical findings are not mutually consistent (Harvey et al., 2019; Rabosky, 2016).

A negative relationship between speciation rate and species range has been rediscovered several times (Jablonski and Roy, 2003; Wagner and Erwin, 1995), but again this pattern has ambiguous interpretations – it might be a consequence of speciation dividing ranges and limiting similarity preventing recolonization, rather than an intrinsic property of species with small ranges.

It is therefore important to come up with models that provide predictions on the relationship between range sizes and speciation rates, and with a conceptual framework providing null expectations on how population size, fragmentation and differentiation co-vary and evolve; how they promote or impede speciation; and how they are transmitted to daughter species.

#### 3.3.1 The freezing voter model

This new archetypal model of speciation takes into account very few factors: migration, mutation and genetic incompatibilities. It can be interpreted as a multi-locus Bateson-Dobzhansky-Muller model where double heterozygotes are unviable, in a rare mutation-rare migration regime, under the infinite-allele model.

As in Section 3.2, this model considers *N* demes forming a graph with edges indicating that migration is possible. Natural choices of graphs include the one-dimensional path (as in the steppingstone model), for mathematical tractability; and the two-dimensional square grid. But other geometries are possible (for instance, a tree could represent a network of rivers or valleys).

Each deme is occupied by a monomorphic population exchanging migrants with a neighboring deme, having then an opportunity to propagate their type inside the target population. However, the migrant is only accepted if her type is sufficiently close, i.e., at a distance smaller than a given threshold, to the type found in the target population, similarly as in the previous section. Finally, mutations occur as in the infinite-allele model.

To be more specific, the model is driven by two types of events:

- **Migration**: at some fixed rate per deme, an individual of the deme migrates towards one of its neighboring populations. The host population then changes type and adopts the type of the migrant, provided the distance between the two types is less than or equal to some fixed threshold.
- **Mutation**: at some fixed rate per deme, a mutation occurs inside the population of the deme and fixes: the whole population changes type to a new type that has never been seen before.

The distance between two types is equal to the number of mutations needed to go from one type to the other (i.e., it is the distance in the genealogy of types, see Section 1.3).

We refer to Figure 4 for a schematic illustration of the dynamics of the model, and to Figure 5 for a computer simulation. In this simulation and throughout the rest of the section, we use a threshold equal to 1.

**Figure 4:**
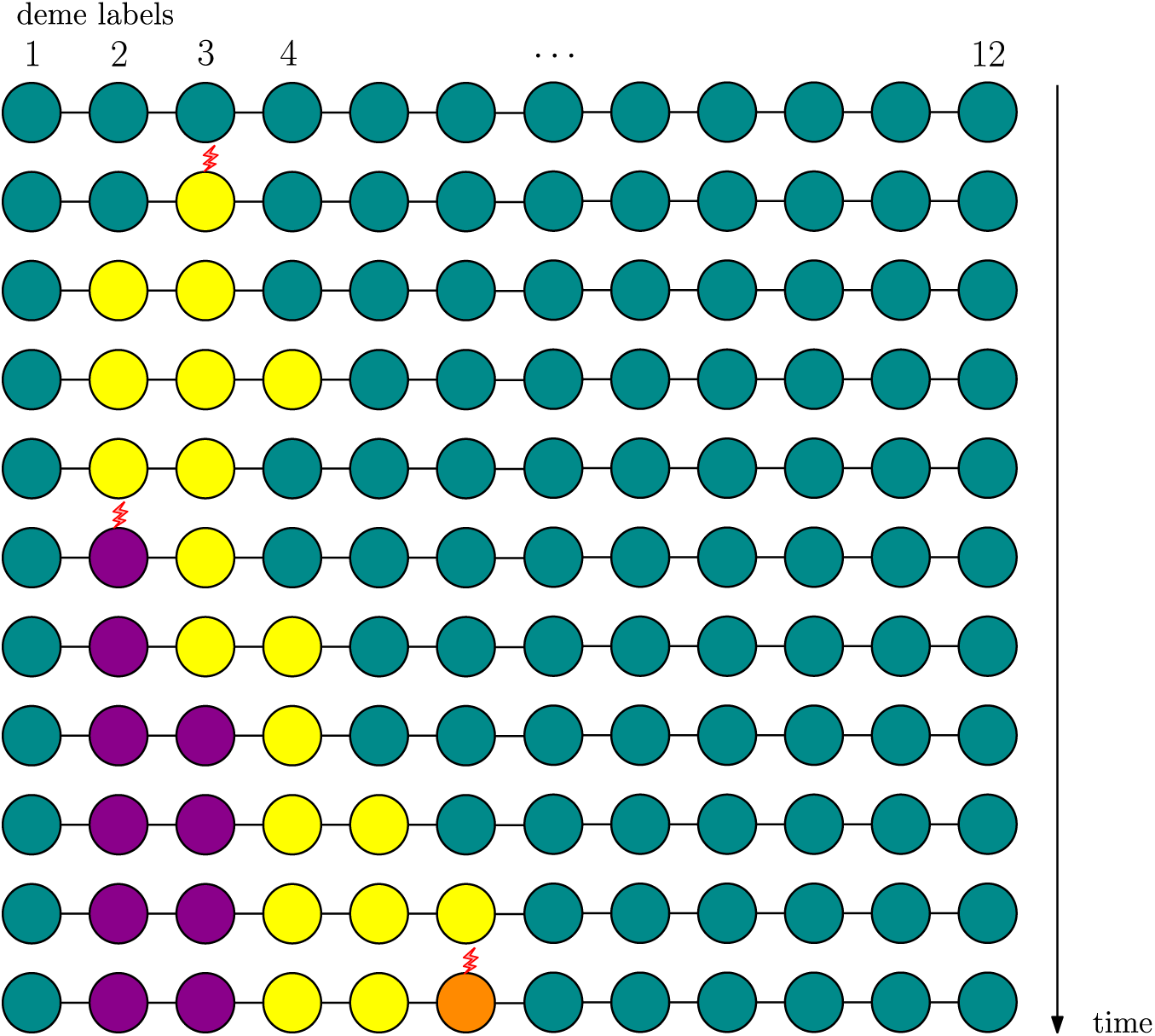
A representation of the freezing voter model, for *N* = 12. Each color represents a type. Each circle is a different deme. Red lightnings indicate mutations. Note that this is a continuoustime model, but only the times where mutations or migrations happen are represented.

**Figure 5:**
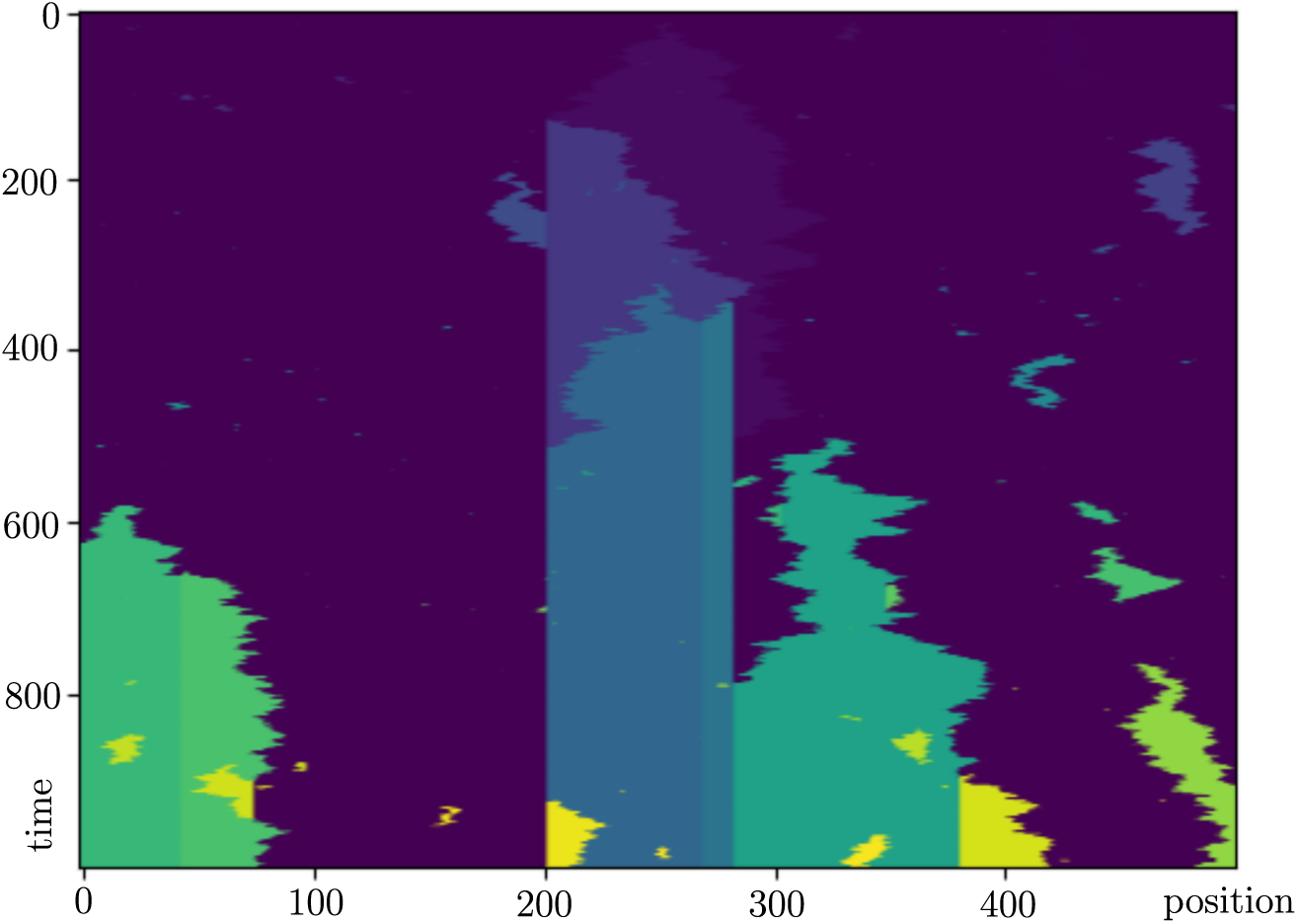
A simulation of the freezing voter model. The horizontal axis represents space. The vertical axis represents time. Each color represents a different genotype. Multiple breeding barriers are formed (vertical lines separating two colors), some of which are temporary (e.g, the barrier between yellow and purple in the bottom left corner) and some of which are permanent.

We will be particularly interested in the formation of what we will call here “breeding barriers”, that is to say of points in space that cannot be crossed by reproductively successful migrants, because of the large genetic distance between populations separated by these points.

As can be seen in Figure 5, some breeding barriers between genetically distant populations may appear and disappear further in time due to the extinction of one of the two types for the benefit of an “intermediary” type that can interbreed with the other populations. Speciation can be defined here as the formation of a *permanent* barrier between two demes, that is to say a point in space that will never be crossed by any successful migration due to the threshold condition on genetic distances.

Thinking of types as political opinions, the model can also be interpreted as describing the dynamics of connected individuals sharing their opinions with their neighbors. Individuals sometimes invent new opinions (mutation) and may succeed in convincing their neighbor (migration), unless the two opinions are too different (breeding barrier). In the case where we start with a fixed number of opinions, no new opinions are formed and there are no restrictions on the propagation of opinions, this is the classic *voter model* introduced by Holley and Liggett (1975) – hence the name “freezing voter model” for our model.

This model provides a rigorous framework to study questions such as

- How long does it take for a “breeding barrier” to appear?
- Where do “breeding barriers” appear?
- What is the typical size and shape of a species range? What is the influence of this shape and of this size on speciation rate?

Though mathematically well-posed, these questions are very challenging to study for the freezing voter model. However, they become easier for a variant of this model, which we now turn to.

#### 3.3.2 A simplification: the Red Queen model

A natural variant of the freezing voter model consists in relaxing the assumption of functional equivalence between species. This can be done in a parsimonious way by introducing a simple asymmetry in gene flow by giving an advantage to recent alleles compared to older alleles in the exchange of genes between interbreeding populations. See Figure 6 for an illustration. A similar hypothesis was made by O’Dwyer and Chisholm (2014), who made an analogy to the Red Queen hypothesis.

**Figure 6:**
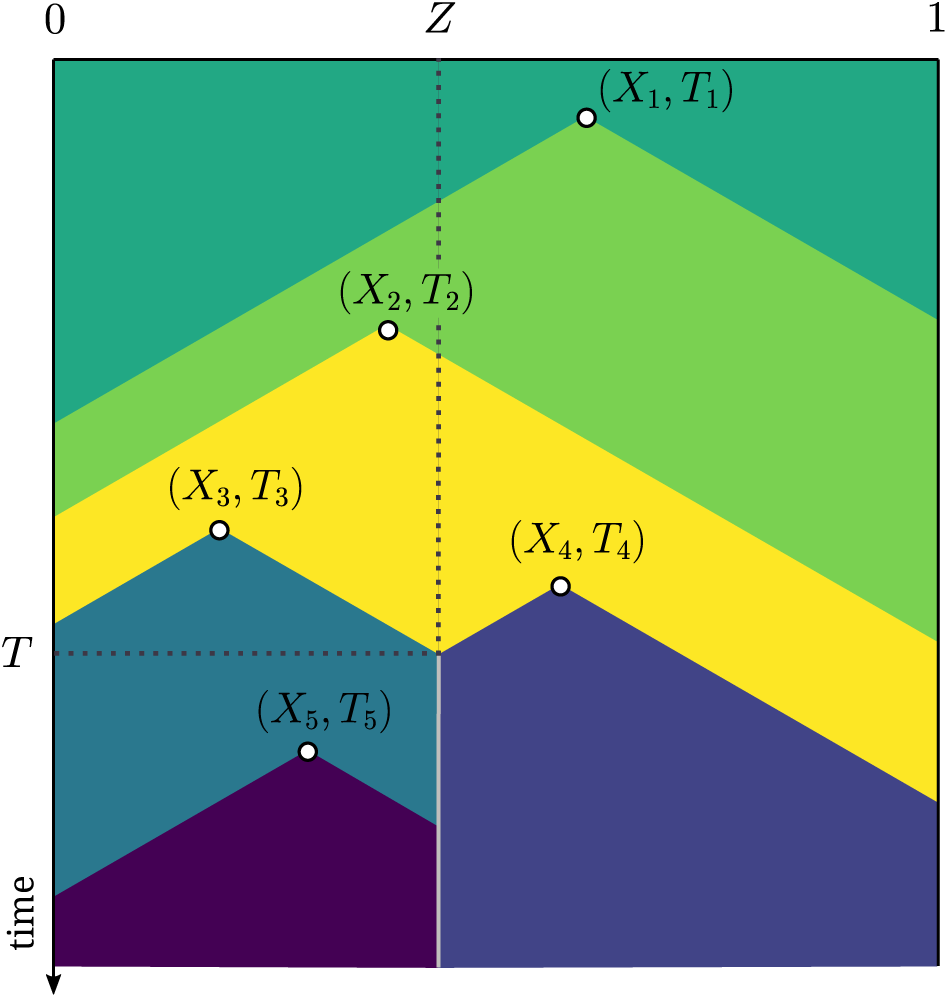
An illustration of the continuous-space limit of the Red Queen model. Time is represented on the vertical axis, and space on the horizontal axis. The (*X_i_, T_i_*) are the positions in space and time of the mutants’ appearances. Each color represents a type. Here, a barrier is formed at time *T* and position *Z*.

In the continuous-space limit obtained by taking the number of demes to infinity, the range of a population carrying a novel type expands at constant speed until it meets the range of a population at genetic distance greater than some threshold value. More precisely, mutants appear at rate *µ* at a uniform position on the interval [0,1]; each mutant propagates to its left and right at speed 1 until it meets another mutant whose genotype is at distance more than 1 from its own genotype, at which point a barrier is formed (speciation event). Ralph and Coop (2010) model parallel local adaptations using similar ideas. In their model, adaptive mutant populations move in waves through continuous space but fronts are not entirely impermeable – as for overlapping tension zones in speciation theory (Barton and Hewitt, 1985, 1989).

In dimension 1 and under the rare-mutation regime (*µ* → 0), we are able to find the distribution of the first barrier that forms between two species, that is to say, the location *Z* of the boundary formed by the first speciation event, and the time *T* at which it arises. Specifically, for small *µ*, the speciation time *T* is well-approximated by an exponential random variable with parameter *^µ^*, and the position *Z* is well-approximated by a Beta(2, 2) variable, i.e. a random variable on [0, 1] with density function *f* (*z*) = 6*z* (1 − *z*) (see Appendix A for a precise statement and a proof of these results). The shape of the species tree can then be derived from computing these distributions and has a *β* index equal to 1 (see Box 4). It is interesting to notice that speciation events are more likely to occur in the center of the range rather than close to its edges, an interesting ecological property known as the “mid-domain effect” (Colwell and Lees, 2000).

## Conclusion

We have set out to demonstrate the advantages of what we termed “archetypal” models of speciation – i.e. models that are built from mechanistic principles and have few parameters. Indeed, macroscopic models and the so-called “lineage-based” approach to diversification have proved very powerful when it comes to portraying macroevolutionary processes; but they are phenomenological in nature and can only shed limited light on the inner causes of macroevolution. By contrast, microscopic models – although typically more challenging to study and harder to relate to realworld data – can, under the right circumstances, yield macroscopic predictions that can be used to assess the validity of our current understanding of evolutionary processes. There is, of course, a trade-off between tractability and realism with this approach, which entails that the goal of an archetypal model can never be to give an accurate description of the evolutionary history of a given taxon, but must instead be restricted to studying specific phenomena in isolation and testing precise hypotheses.

We have reviewed some of the main existing archetypal models of speciation, in order to give an overview of their diversity and of the questions that they can address, but also to point out some of their current limitations.

The models we have presented are either built from mechanistic constructions (e.g. the Higgs– Derrida model and related variants) or more heuristic ones (e.g. Hubbell’s point-mutation model, protracted speciation models). In any case they all aim at explaining the main patterns of speciation (e.g. species abundance distributions, shapes of phylogenies, species-area relationships) by representing possibly complex evolutionary processes using only a few effective parameters – e.g. speciation rates, ecological community sizes, mutation and/or migration rates… We have categorized archetypal models into three classes: **(1) clonal evolution models**, **(2) models of genetic isolation**, and **(3) models of isolation by distance**. For each of these three classes, the main process responsible for homogenizing gene pools is **(1) ecological drift**, **(2) gene flow**, and (3) **spatial drift**. Of course, many models mix these three types of homogenizing processes; but most models put the emphasis on one of these barriers to differentiation, with various degrees of purity:

1. **Clonal evolution models** assume that lineages diverge only through parallel accumulation of mutations. In turn, speciation can only be slowed down by stochastic birth and death events purging this diversity. These models include: the point-mutation model of speciation (Hubbell, 2001; Chave, 2004; Jabot and Chave, 2009); the protracted speciation model (Rosindell et al., 2010; Etienne and Rosindell, 2012; Lambert et al., 2015); and the model of speciation by genetic differentiation (Manceau et al., 2015).
2. **Models of genetic isolation** specifically ask whether pairs of individuals or populations are genetically compatible. Here, various processes can contribute to homogenizing gene pools: sexual reproduction recombines genomes and can hence purge some alleles, migration between demes enables gene flow and the spread of local diversity. Furthermore, the rates of homogenizing processes may decrease with increasing dissimilarity. These models include the Higgs–Derrida model (Higgs and Derrida, 1991, 1992); the split-and-drift random graph (Bienvenu et al., 2019); the parapatric model of Gavrilets (2000) and its extensions (Yamaguchi and Iwasa, 2013, 2015; Miró Pina and Schertzer, 2019); and the gene-based diversification model (Marin et al., 2020).
3. **Models of isolation by distance.** Homogenization here is mainly controlled by dispersal, range expansion or local extinction. These models include the “extinction-recolonization” model of Gavrilets et al. (2000a), the “topopatric” model of de Aguiar et al. (2009) and the geographic model of speciation of Pigot et al. (2010).

We have identified a set of key mechanisms that are rarely taken into account by these models (the failure of speciation at secondary contact, the feedback of dissimilarity on homogenization, the emergence in space of reproductive barriers), and we have proposed new models incorporating them. These new models open up promising avenues of research – both for the mathematical challenges that they pose and, more importantly, for the insights into evolutionary processes that their understanding will provide.

A crucial aspect of the micro-macro approach – which we have already mentioned, but which is beyond the scope of this article – is the comparison with real-world data. Indeed, microscopic models are designed based on our current knowledge of evolutionary processes, and in a sense they are the mathematical formulation of our understanding of these processes. Their design and study is an interesting scientific topic in itself; but it is not an end-goal. Instead, the end-goal is to infer their validity from real data, in order to see whether our current understanding of speciation is compatible with reality – and update it accordingly. As of today, there is no consistent relation between the predictions of microscopic models and macroscopic data on the topic of speciation. This reflects the fact that the main processes driving speciation and the interplay between these processes are not yet fully understood.

## Acknowledgments

FB was supported by Dr. Max Rössler, the Walter Haefner Foundation and the ETH Zürich Foundation. AL thanks the *Center for Interdisciplinary Research in Biology* (CIRB, Collège de France) and the *Institute of Biology of École Normale Supérieure* (IBENS, École Normale Supérieure, Université PSL) for funding. The authors thank Félix Foutel-Rodier, Yannic Wenzel and Philibert Courau for their comments and enlightening discussions. We also wish to thank a Guest Editor (Théo Gaboriau) and two anonymous reviewers for their very thorough reading of the first version of this work and the numerous improvements that they suggested.

## Appendices

### A Appendix on the Red Queen model

In this appendix, we study the position and time of apparition of the first speciation event in the Red Queen model of Section 3.3.2. The goal is to give a sketch of the proof of the following theorem focusing on the important ideas but leaving aside uninformative technicalities.

Theorem 1. Let Z and T be, respectively, the position and time of the first speciation event in the continuous-space limit of the Red Queen model with mutation parameter µ. Then, as µ goes to 0, (*Z, µ*^2^*T*) *converges in distribution to* (*Z*_lim_, *T*_lim_), *where*

i. Z_lim_ a continuous random variable on [0, 1] with probability density function f (z) = 6z (1 − z);
ii. T_lim_ is an exponential random variable with parameter 1/3;
iii. *(iii) Z*_lim_ *and T*_lim_ *are independent*.

Note that, in this theorem as in the rest of this section, the random variables *Z* and *T* depend on the parameter *µ*, but that we keep this dependence implicit for readability. We will also do so for other quantities.

Finally, throughout the rest of the section, we use the word *barrier* to refer to a point in [0, 1] that cannot be crossed after a speciation event (in other words, the place where to two mutants involved in the speciation event meet); see Figure 6.

### A.1 Proof idea: limit of *T*

Let us start by introducing some notation. Let *P* = (*T_i_, U_i_*) *_i_*_∈N_∗ be the set of times and positions of apparition of the mutants, where *T_i_* is the time of apparition of the *i*-th mutant and *U_i_* is the point in [0, 1] where it appeared. Note that *P* is a Poisson point process on R_+_ × [0, 1] with intensity measure *µ*d*t* ⊗d*x*. Thus, (*U_i_*)*_i_*_∈N_∗ is a sequence of independent and identically distributed (i.i.d.) uniform random variables on [0, 1], and there exists a sequence (*ξ_i_*)*_i_*_∈N_∗ of i.i.d. exponential variables with parameter *µ* such that (*ξ_i_*)*_i_*_∈N_∗ is independent of (*U_i_*)*_i_*_∈N_∗ and, for all *i* ∈ N^∗^,

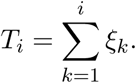

Each mutant that appears at position *x* and time *t* will propagate to the left and to the right at speed 1. Therefore, it will encounter a younger mutant (i.e. an individual carrying a genotype that appeared after *t*) if and only if a mutant appears in the set *A_x,t_* corresponding to the yellow zone in Figure 7. Formally, for all *t* ∈ R_+_, *x* ∈ [0, 1], let

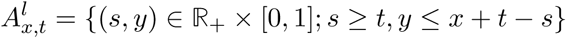

and

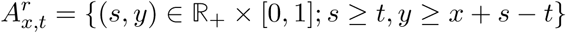

correspond to the left and right components of this set. Then, *A_x,t_* = *A^l^_r x,t_*. The area of *A_x,t_*, which we denote by *λ*(*A_x,t_*), is

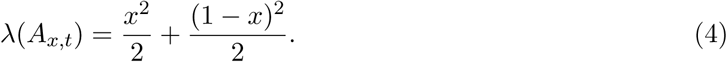

**Figure 7:**
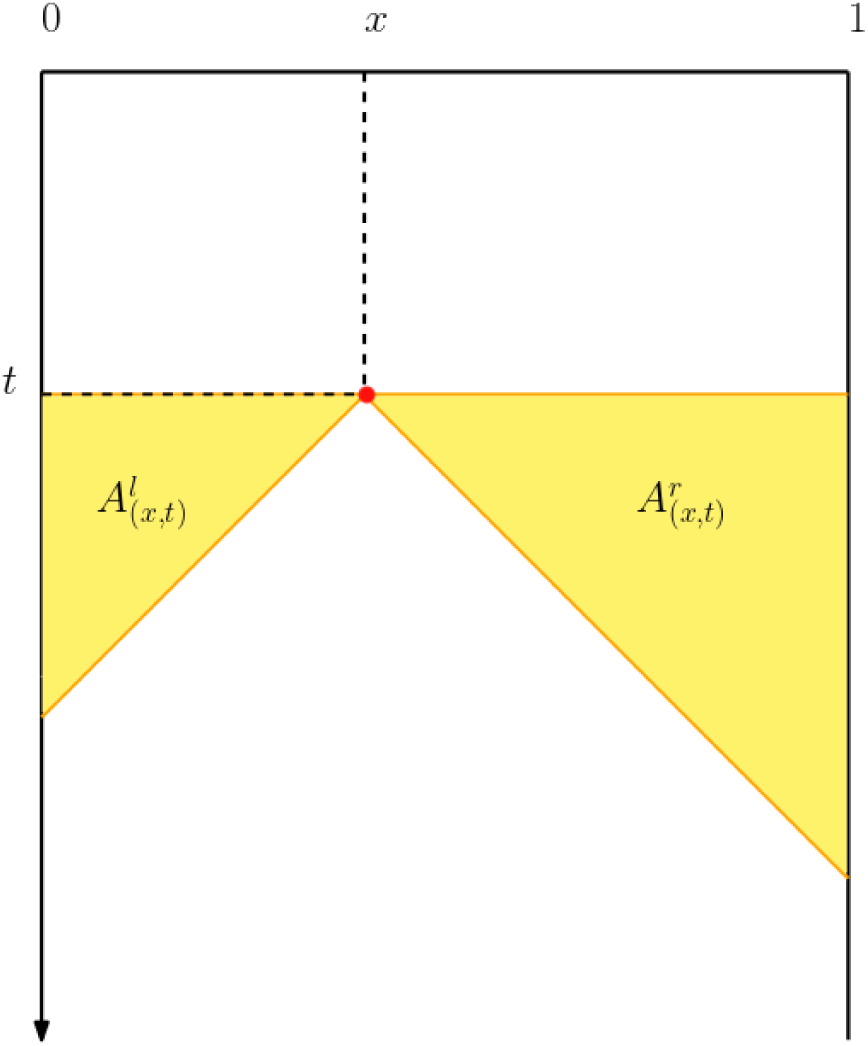
The mutation represented by a red point at (*t, x*) will create a barrier with a mutant that appeared after *t* if and only if at least one mutant appears in the yellow zone.

So far, we have assumed that the position where the first mutant appears is fixed, but in reality it is a random variable (*X, θ*) ∈ [0, 1] × R_+_, where *X* is uniform on [0, 1]. Therefore, *λ*(*A_x,t_*) is the area of the zone where the apparition of a new mutant will create a barrier, “conditional on (*X, θ*) = (*x, t*)” (here we use quotes because the probability that (*X, θ*) = (*x, t*) is equal to 0 for all (*x, t*), but this quantity is well-defined as a conditional expectation). Integrating against the law of *X*, we get that the (unconditional) expected area of this zone is

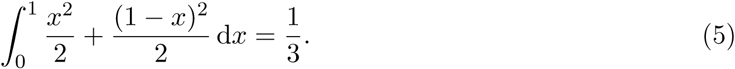

Therefore, if we denote by *I* the index of the first mutant involved in a speciation event, and by *T_I_* the time of apparition of that mutant, since all mutant appear at rate *µ* we have

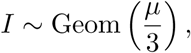

i.e. *I* follows a geometric distribution with parameter 1*/*3. Moreover, since the (*ξ_i_*)*_i_*_≥1_ are exponentially distributed with parameter *µ*, an easy computation yields

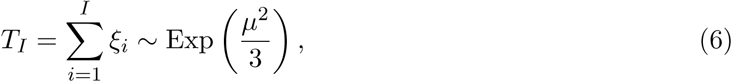

where Exp(*θ*) denotes the exponential distribution with parameter *θ* and ~ indicates equality in distribution.

Recall that *T* is the time of the first speciation event, that is to say the time when the *I*-th mutant and the other mutant involved in this speciation meet (in fact, as explained below, that other mutant is the (*I* +1)-th mutant with arbitrarily large probability as *µ* goes to 0; but that is not relevant for now). Since all mutants propagate at speed 1 in the interval [0, 1], once the *I*-th mutant has appeared it cannot take more than one unit of time for the speciation to occur, irrespective of the initial positions of that mutant and of the other mutant involved in the speciation.

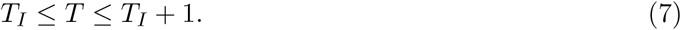

Therefore, Combining Equations (6) and (7) gives that, as *µ* goes to 0, *µ*^2^*T* converges in distribution to an exponential random variable with parameter 1*/*3, proving point (i) of the theorem.

**Figure 8:**
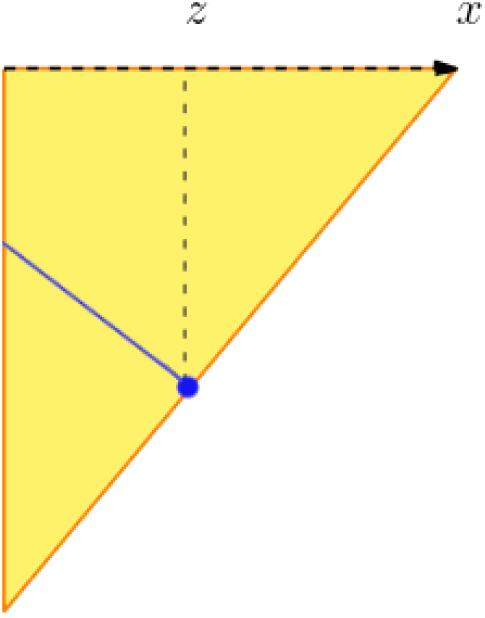
The blue line represents the set of points such that the apparition of a second mutant at one of these points will create a barrier at position *z*.

### A.2 Proof idea: limit of *Z*

Keeping the notation of the previous section, recall that *I* is the index of apparition the first mutant involved in the first speciation event. When the mutation rate goes to 0, the probability that three mutants coexist between the apparition of the *I*-th mutant and the first speciation goes to 0. This entails that the other mutant involved in the first speciation event will have index *I* + 1 with probability that goes to 1 as *µ* goes to 0. Therefore, in the limit, the position of the first barrier is the point where the mutants *I* and *I* + 1 meet.

Note that if the *I*-th mutation appears at *x* ∈ [0, 1], then the *I*-th and (*I* + 1)-th mutants meet at some point *z* ∈ [0, *x*] if and only if the (*I* + 1)-th mutant appears somewhere in the blue segment depicted in Figure 8. The length *L^l^* of this segment is

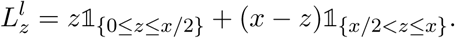

Similarly, the set of potential points of apparition of the mutant *I* +1 that yield a barrier formed at *z* ∈]*x*, 1] has length

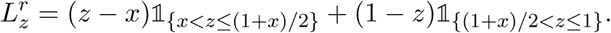

Thus, for any *z* ∈ [0, 1], the probability density of *Z*_lim_ in *z* is proportional to *L_z_* = *L^l^* + *L^r^*.

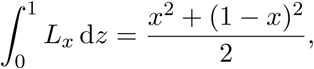

Since we get that, conditional on {*X_I_* = *x*}, *Z* has density

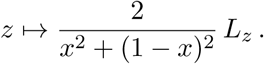

To finish the proof, it suffices to determine the law of *X_I_* and to integrate the conditional density of *Z* against it. The probability density of the position of apparition of a mutation that generates a speciation at position *x* is proportional to the area of the yellow zone in Figure 7, that is to say 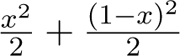. Therefore, the probability density of *X* is

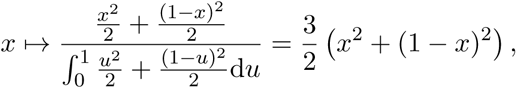

and by integrating the conditional density of *Z*_lim_ against this density, we get that *Z*_lim_ has density *z* I→ 6*z*(1 − *z*), concluding the proof.

